# Dynamic visualization of high-dimensional data

**DOI:** 10.1101/2022.05.27.493785

**Authors:** Eric D. Sun, Rong Ma, James Zou

## Abstract

Dimensionality reduction (DR) is commonly used to project highdimensional data into lower dimensions for visualization, which could then generate new insights and hypotheses. However, DR algorithms necessarily introduce distortions in the visualization and cannot faithfully represent all relations in the data. As such, there is a need for methods to assess the reliability of DR visualizations. Here we present DynamicViz, a framework for generating dynamic visualizations that capture the sensitivity of DR visualizations to perturbations in the data. DynamicVic can be applied to all commonly used DR methods. We show the utility of dynamic visualizations in diagnosing common interpretative pitfalls of static visualizations and extending existing single-cell analyses. We introduce the variance score to quantify the dynamic variability of observations in these visualizations. The variance score characterizes natural variability in the data and can be used to optimize DR algorithm implementations. We have made DynamicViz freely available to assist with the evaluation of DR visualizations.

## 1 Introduction

Dimensionality reduction (DR) is a standard practice in the analysis of highdimensional data. In addition to reducing noise and facilitating downstream computational analyses, DR methods are widely used to visualize data in two or three dimensions. In fact, many DR methods have been expressly developed for generating informative visualizations of high-dimensional data. Some popular DR methods for data visualization include the linear principal component analysis (PCA) and the nonlinear t-distributed stochastic neighbor embedding (t-SNE) [1] and uniform manifold approximation and projection (UMAP) [2]. Many other DR methods have been developed to address shortcomings of the commonly used t-SNE and UMAP approaches [3–5]. DR methods for visualization have found specific uses across a wide range of different disciplines. Some examples include validating cell type identities in single-cell biology [6, 7], probing input embeddings from deep learning models, exploring geographic patterns in human genomics [8], and dissecting chemical abundance of stellar objects [9].

Despite the popularity of DR methods for visualizing high-dimensional data, these methods are prone to distortions and heterogeneity in the quality of the low-dimensional visualization [6, 10–14]. As such, naive use of DR methods to validate, confirm, or inform research findings and directions can be susceptible to misinterpretation due to these distortions. For example, in the field of single-cell biology, t-SNE or UMAP visualizations are often used to confirm cell type identities of clusters [6], integrate different single-cell datasets [15– 17], and compute cell trajectories using RNA velocity measures [18, 19]. For each of the aforementioned use cases, distortions in distances between observations and heterogeneities in the quality of the DR visualization are present and can affect the resulting interpretations [10, 13, 20–22]. Generally, through these distortions, DR visualizations may lead to improper validation of clusters (i.e. under-clustered or over-clustered), artificial detection or lack of detection of bridging connections between clusters, and artificial presence of ordering or loss of ordering of observations along metadata axes.

The concerns of DR are exacerbated by the static nature of current DR visualization methods, which typically show an image representing a single initialization of the DR method. This approach can mask randomness in the visualization due to the data and/or the DR method, and can be vulnerable to cherry-picking. To address these limitations, we introduce DynamicViz, a framework for generating dynamic visualizations of data by aligning multiple bootstrapped DR visualizations. As a result, dynamic visualization provides strictly greater information than any single static visualization. These dynamic visualizations provide a visually intuitive framework for users to understand the sensitivity of DR visualization to perturbations in the data and to any stochastic components of the DR method. We show that this tool can be used to diagnose a variety of interpretative pitfalls, which encompass many of the aforementioned issues. To quantify the degree of fluctuation of a sample across the bootstrap DR visualizations, we introduce the variance score, which tracks with variability across real-world replicates and can be used to optimize the DR visualization pipeline. Unlike previous quality metrics for assessing DR visualizations that measure concordance of the visualization with the high-dimensional data [11, 12, 14, 23–26], the variance score can measure the effect of sampling noise in data generation on the distortions observed in DR visualization. In order to improve the accessibility of tools for evaluating DR visualizations, we have released DynamicViz as an open-source Python package that includes tools for generating dynamic visualizations of data and calculating the variance score among other functionalities.

## 2 Results

Dynamic visualization of high-dimensional data aims to provide strictly more visual information than what is available in existing static DR visualizations. The inputs for dynamic visualization and assessment are the same as those for the chosen static DR method with the exception of one additional parameter corresponding to the number of bootstrap samples used for the visualization, which is dependent on the intended use case and computational resources available to the user. Dynamic visualizations can provide robust confirmation of prior findings and generation of new research hypotheses.

### 2.1 Dynamic visualization procedure

We represent the input data for dynamic visualization as *X* ∈ ℝ^*n*×*p*^, where *n* is number of observations and *p* is the number of features. For example, in the case of single-cell transcriptomics data, the input data is a normalized transcript count matrix where observations correspond to individual cells and the features correspond to a typically large number of genes. Next, we generate a bootstrap sample of *X* by sampling *n* rows of *X* with replacement. This is repeated *B* times to accumulate a set of *B* bootstrap samples of *X*, which we denote as {*X*^(1)^, …, *X*^(*B*)^ }. For a given DR method *f* (·), a two-dimensional visualization of *X* is generated, which we refer to as *Y* = *f* (*X*) ∈ ℝ^*n*×2^. Using the same DR method, visualizations are also generated for the bootstrap samples, yielding the set {*Y* ^(1)^, …, *Y* ^(*B*)^} in a random order. Each bootstrap visualization *Y* ^(*k*)^ with *k* ∈ {1, …, *B*} is aligned to the reference visualization *Y* using the Kabsch algorithm to minimize the root mean square distance between concordant observations of *Y* ^(*k*)^ and *Y* under three-dimensional rotation operations, which preserve distance relations in the visualization and thus do not change the value of the objective function minimized by most DR methods (see Methods for details).

Finally, the sequence of randomly ordered bootstrap visualizations {*Y* ^(1)^, …, *Y* ^(*B*)^}is presented dynamically. Currently, we provide three different dynamic visualization modalities, each highlighting some aspects of the data and its DR visualizations. Interactive visualization using HTML produces interpolated transitions between the random sequence of bootstrap visualizations and provides users with high-level interactions such as toggling observations on/off by discrete labels and reading metadata embedded in the visualization. Animated visualization presents a random sequence of bootstrap visualizations in video and GIF formats, which can be easily embedded in research workflows and presentations. Stacked visualization overlays the aligned bootstrap visualizations in a single image with low opacity of scatter points to visually represent density of visualizations, which is well-suited for presenting the dynamics in a single frame. An illustration of the dynamic visualization procedure is shown in Figure 1A and examples of each mode of dynamic visualization are shown in Figure 1B-G. Example interactive visualization HTML files and animated visualization GIF files corresponding to Figure 1 are available in Supplementary Videos 1-4.

**Fig. 1.**
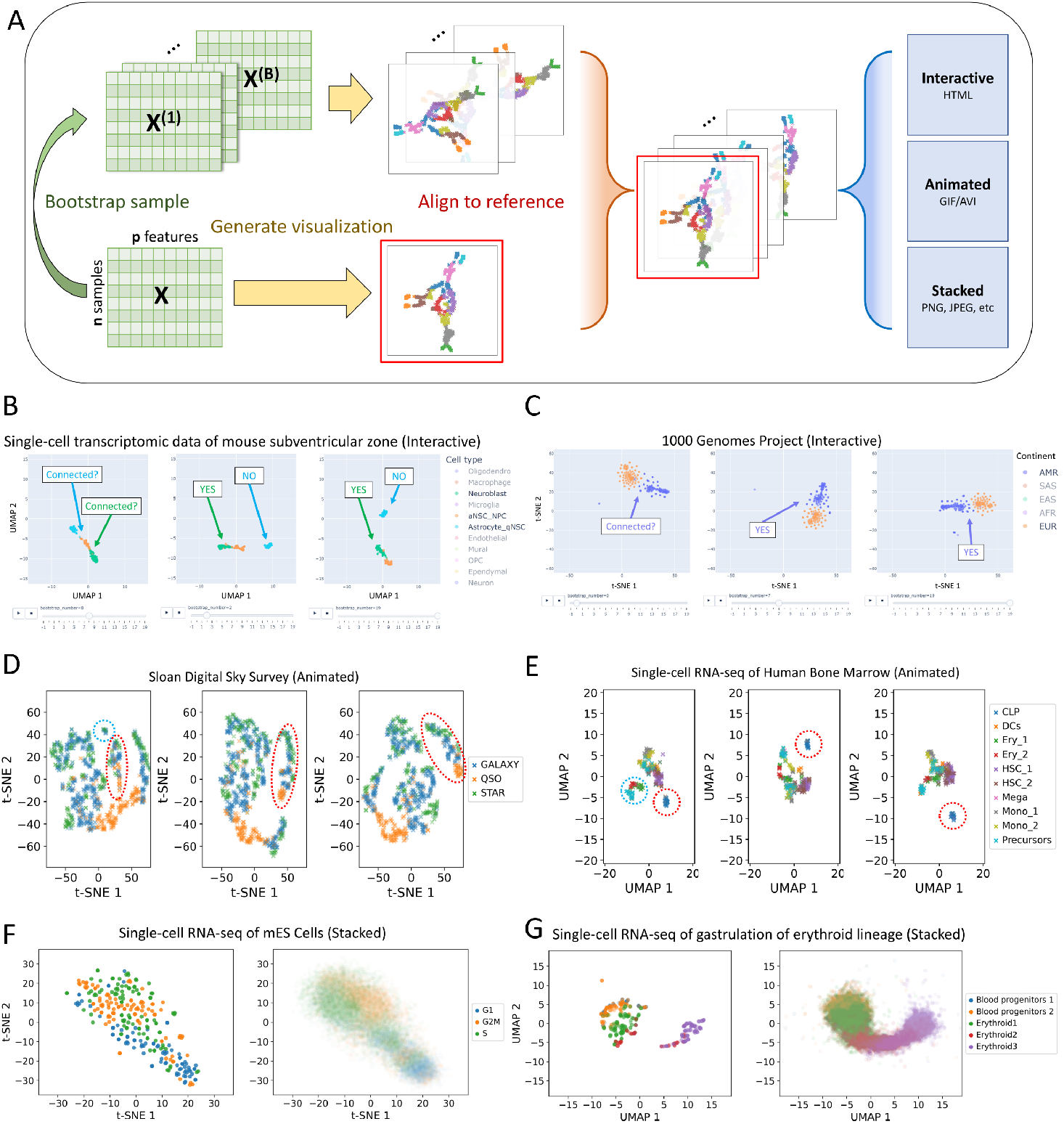
Dynamic visualization and its uses. (A) Schematic illustration of the dynamic visualization pipeline with bootstrapping of the data, generating DR visualizations, rigid alignment of the bootstrapped visualizations and final presentation with either interactive, animated, or stacked modalities. (B) Interactive visualization of single-cell transcriptomic data from mouse subventricular zone for diagnosing stability of bridging connections between cell type clusters in the neural stem cell lineage. (C) Interactive visualization of genomic data from the 1000 Genomes Project for diagnosing stability of bridging connections between European and Ad-Mixed American population clusters. (D) Animated visualization of spectral data of stars, quasars, and galaxies from the Sloan Digital Sky Survey queries stability of clusters identified in one visualization (leftmost panel) across other visualizations (consistent clusters are circled in red, inconsistent clusters are circles in blue). (E) Animated visualization of single-cell transcriptomic data from human bone marrow queries stability of clusters identified in one visualization (leftmost panel) across other visualizations (consistent clusters are circled in red, inconsistent clusters are circles in blue). (F) Stacked visualization of transcriptomic data of mouse embryonic stem cells undergoing three different phases of cell cycle (right) reveals stable separation of cell cycle phases that is not apparent from a single visualization (left). (G) Stacked visualization of single-cell transcriptomic data from gastrulation of erythroid lineage (right) reveals stable order of cell types along a trajectory that is not apparent from a single visualization (left).

### 2.2 Dynamic visualization diagnoses stability of bridging connections between clusters

Bridging connections often occur in the visualization of clustered data where two or more distinct clusters of observations may appear to be in contact or have a narrow connection between them in the low-dimensional visualization. These bridging connections may represent underlying relationships between the two clusters, particularly in single-cell biology where bridging connections may capture cell differentiation or developmental pathways, but they may also be artifacts created by distortions in the DR method and/or intrinsic noise in the data. The validity of a bridging connection is difficult to discern from a static visualization.

Dynamic visualizations can help to discriminate robust bridging connections that appear across most bootstrap visualizations from incidental or artificial bridging connections that only appear in one or a small minority of bootstrap visualizations. For example, interactive UMAP visualization of single-cell transcriptomes of the mouse subventricular zone [27], a neurogenic region in the brain, highlights a robust bridging connection between the activated neural stem cell / neural progenitor cell (aNSC-NPC) cluster and neuroblast cluster, and a much less stable connection between the aNSC-NPC and astrocyte cluster (see Figure 1B). From a single visualization, the relative stability of these connections is not apparent (see leftmost frame of Figure 1B). Across UMAP visualizations of four independent mouse SVZ single-cell transcriptomic datasets, bridging connections between aNSC-NPC and neuroblast clusters were present in all visualizations and connections between aNSC-NPC and astrocyte clusters were present in only two visualizations, consistent with our findings from dynamic visualization. In a second example, interactive t-SNE visualization of human genomics data from the 1000 Genomes Project [28, 29] reveals a stable bridging connection between genome clusters corresponding to admixed American (AMR) and European (EUR) populations (see Figure 1C). In both cases, dynamic visualization is useful in assessing the relative stability of bridging connections between clusters.

### 2.3 Dynamic visualization assesses confidence in cluster hypotheses

DR visualizations of clustered data are often used to either confirm the labels obtained from a clustering algorithm or identify subsets of observations that may represent new clusters. Dynamic visualizations can be used in both cases to study the stability of identified clusters across different bootstrap visualizations. Animated t-SNE visualization of galaxy, quasar, and star spectra data from the Sloan Sky Digital Survey [30] reveals potential clusters and their consistency across different bootstrap visualizations, which suggests that the larger of two clusters identified in one frame is stable while the smaller cluster is not (Figure 1D). Of note, the larger cluster has mixed labels from all three astronomical classes, suggesting that further characterization of this cluster may yield insight into the distinguishing properties of its members. Likewise, animated UMAP visualization of human bone marrow single-cell transcriptomic data [31] shows several potential disjoint clusters with only the clonal progenitor cell (CLP) cluster consistently present across most bootstrap visualizations (Figure 1E), thus supporting the identification of the CLP cluster.

### 2.4 Dynamic visualization reveals separation and ordering of data by label

In both clustered and non-clustered data, visual patterns in observation labels are useful for greater understanding of relations between the labels and for generation of new research hypotheses. Dynamic visualizations can be used to assess subtle patterns in DR visualizations such as degree of separation of different labels in a non-clustered setting and ordering of labels along a continuous trajectory. For example, static t-SNE visualization of gene expression data from mouse embryonic stem cells [32] does not reveal clear separation of cells under the label of cell cycle phase while dynamic t-SNE visualization through the stacked modality shows enhanced separation of the three cell cycle phase labels (Figure 1F). Static UMAP visualization of single-cell transcriptomic data from gastrulation of the erythroid lineage [33] show disjointed clusters with no apparent differentiation trajectory while dynamic UMAP visualization of the same data reveals a continuous trajectory through biologically relevant cell states (Figure 1G). In both cases, the application of dynamic visualizations recovers the underlying biological truth.

### 2.5 Variance score for quantifying fluctuation in dynamic visualization

To quantify the instability shown in dynamic visualization, we introduce the variance score, which measures the average variance in pair-wise Euclidean distances between a given observation and its neighbors across all bootstrap visualizations. For a set of bootstrap data {*X*^(1)^, …, *X*^(*B*)^} and DR method *f* (·) yielding visualizations {*Y* ^(1)^, …, *Y* ^(*B*)^}, we define the variance score for the *i*-th observation over its neighborhood *N*_*i*_ as:

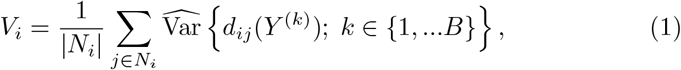

where

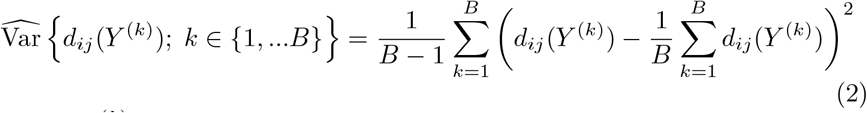

and *d*_*ij*_(*Y* ^(*k*)^) denotes the Euclidean distance between the observation *i* and a neighboring observation *j* in the bootstrap visualization *Y* ^(*k*)^ (see Methods for details on how *d*_*ij*_ is normalized). The *i* and *j* indices are realized with respect to the original data *X* such that *i* and *j* refer to consistent observations across the bootstrap visualizations. An illustration of Eq. (1) is shown in Figure 2A. In our analyses, we used the global neighborhood such that for any observation, its neighborhood consisted of all other observations in the visualization. Additional neighborhood definitions are presented in Methods.

**Fig. 2.**
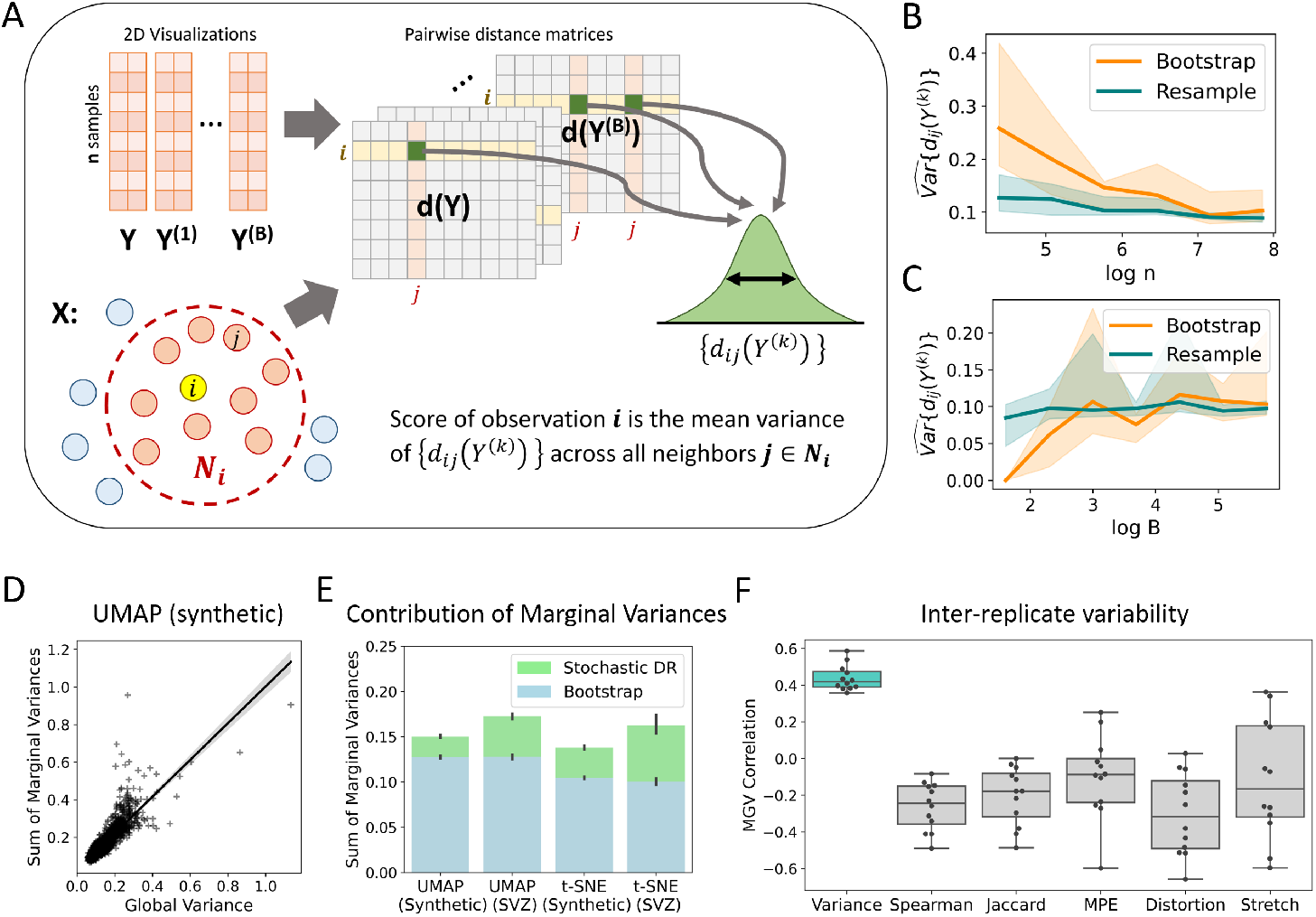
Variance score and its properties. (A) Schematic illustration of variance score calculation from the bootstrapped DR visualizations by computing variance of pairwise distance between any two observations denoted i and j across all bootstrapped visualizations and averaging this variance measure over all neighbors j to obtain a variance score for i. (B) Variance in the Euclidean distance between two predetermined observations in UMAP visualization across either 100 bootstrap sample data or 100 resampled data from a Gaussian mixture model (50 features, 5 distributions) for different numbers of observations, *n*. 95% confidence intervals are shown for 20 pairs of predetermined observations. (C) Same setting as panel C except with *n* = 1000 and different number of bootstrap samples or resamples, *B*. Sum of the two marginal variance scores for UMAP from eliminating DR stochasticity or eliminating bootstrap sampling of data respectively compared to the total variance score for UMAP on high-dimensional data drawn from a mixture of five Gaussian distributions (*n* = 1000, *p* = 50). (E) Relative contributions of the marginal variance scores for t-SNE and UMAP visualizations of synthetic data drawn from a mixture of five Gaussian distributions and of single-cell transcriptomic data from mouse subventricular zone. (F) Spearman correlation between DR quality metrics (including variance score) computed on a single replicate and the mean gene variance across all 12 technical and biological replicates in the MER-FISH mouse primary motor cortex spatial transcriptomics dataset. In panels D-F, variance scores were computed for 100 bootstrap samples.

Bootstrap sampling is widely used in statistics to quantify variance in the data and in the analysis method. By construction, the variance score defined in Eq. (2) is expected to capture the variability of a visualization method when independent replicates are available for a given dataset of interest (see Methods). We show empirical convergence of the variance scores based on bootstrap samples to those based on actual independent replicates, as the number of observations *n* or the number of bootstrap samples *B* increases, demonstrating the interpretation of variance scores as the locally averaged conditional variance 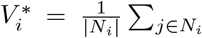 Var(*d*_*ij*_(*f* (*X*))|*X*_*i*_, *X*_*j*_) (see Eq. (7)) in the large sample limit (Figure 2BC and Extended Data Figure A1BC). In addition, we observe similar silhouette scores, a metric of clustering quality, between visualizations generated from repeated samples, each consisting of *n* observations, from a mixture of Gaussian distributions and those generated from bootstrap sampling of one sample from the mixture of Gaussian distributions (Extended Data Figure A1A).

### 2.6 Marginal variance scores highlight two sources of variability in standard DR visualization pipeline

By the law of total variance, Var(*d*_*ij*_(*f* (*X*)) |*X*_*i*_, *X*_*j*_) captures two sources of variability in the final visualization, that is, the variability due to the underlying data distribution, and the variability due to the possibly random visualization method *f*. This relationship can be translated to a decomposition of the variance score

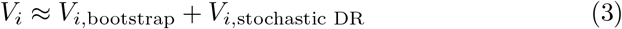

where

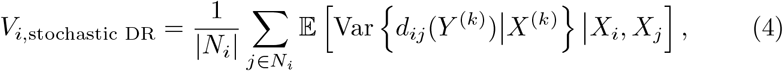

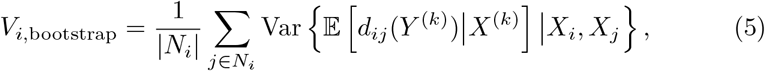

and *X*^(*k*)^ is a random bootstrap sample of *X*. Here, *V*_*i*,stochastic DR_ represents the variance score that is attributable to the stochastic procedure of the DR method *f* (·) and *V*_*i*,bootstrap_ represents the variance score that is attributable to the bootstrap perturbation to the data. In the case where the DR method is non-stochastic, then *V*_*i*,stochastic DR_ = 0 and *V*_*i*_ ≈ *V*_*i*,bootstrap_. For a complete statement of this result and its conditions, refer to Methods.

Empirically, we observe the decomposition of the variance score into these two marginal variance scores in several settings. The sum of the estimated marginal variance scores (average taken across 10 random seeds for DR initialization and 10 random seeds for bootstrap sampling) is approximately equal to the variance score from Eq. (1) for both UMAP and t-SNE visualizations of synthetic data sampled from high-dimensional mixture of Gaussian distributions and single-cell transcriptomic data from mouse SVZ (see Figure 2D). Across these case examples, *V*_*i*,stochastic DR_ is generally smaller than *V*_*i*,bootstrap_ (see Figure 2E). As such, in certain contexts, *V*_*i*_ may be a good approximation for *V*_*i*,bootstrap_ even when the DR algorithm is stochastic.

### 2.7 Variance score tracks with natural variability in data

The primary objective of dynamic visualization and the variance score is to simulate and quantify the fluctuations in DR visualization subject to noise in the data and distortions from the DR algorithm. To test whether the variance score captures variability from experimental sampling noise, we independently computed UMAP variance scores for each of 12 technical and biological replicates in a spatial transcriptomics dataset of mouse primary cortex from the VizGen MERFISH platform [34], which contained 24 cell types. Since there is no concordance between individual cells across the replicates, we calculate a mean variance score at the level of cell type and compare this to the mean gene expression variance (MGV) computed across all 12 replicates for each cell type. The MGV is computed as the variance of the mean gene expression of each gene across all cells of a given cell type. Compared to existing metrics for assessing DR visualization quality, the variance score is consistently correlated with the MGV across all replicates (see Figure 2F).

### 2.8 Using variance scores to optimize choice of DR algorithms

A challenge in the DR visualization of high-dimensional data is to select hyperparameter values for the DR algorithm (e.g. perplexity in the t-SNE algorithm), which is highly dependent on properties of the data. A common approach is to select hyperparameter values that appear to yield convincing visualizations in a heuristic manner [6, 20]. Other approaches have been proposed for selecting hyperparameter values based on optimization of an objective function [14, 35, 36]. Here, we present minimization of the variance score as a natural procedure for selecting the most reliable hyperparameters for DR visualization, which by definition, will yield the least expected variability in DR visualization across replicates of the data and across different random states for the DR algorithm. For illustration, the variance score for t-SNE visualizations of mouse SVZ single-cell transcriptomic data is minimized for an intermediate perplexity value of 160. This optimal perplexity value produced better visual separation and stability of cell type clusters than other choices of perplexity, as shown through stacked representation of all bootstrap visualizations (Figure 3A). Additional examples of optimizing DR visualization hyperparameters using the variance score are shown in Extended Data Figure A2.

**Fig. 3.**
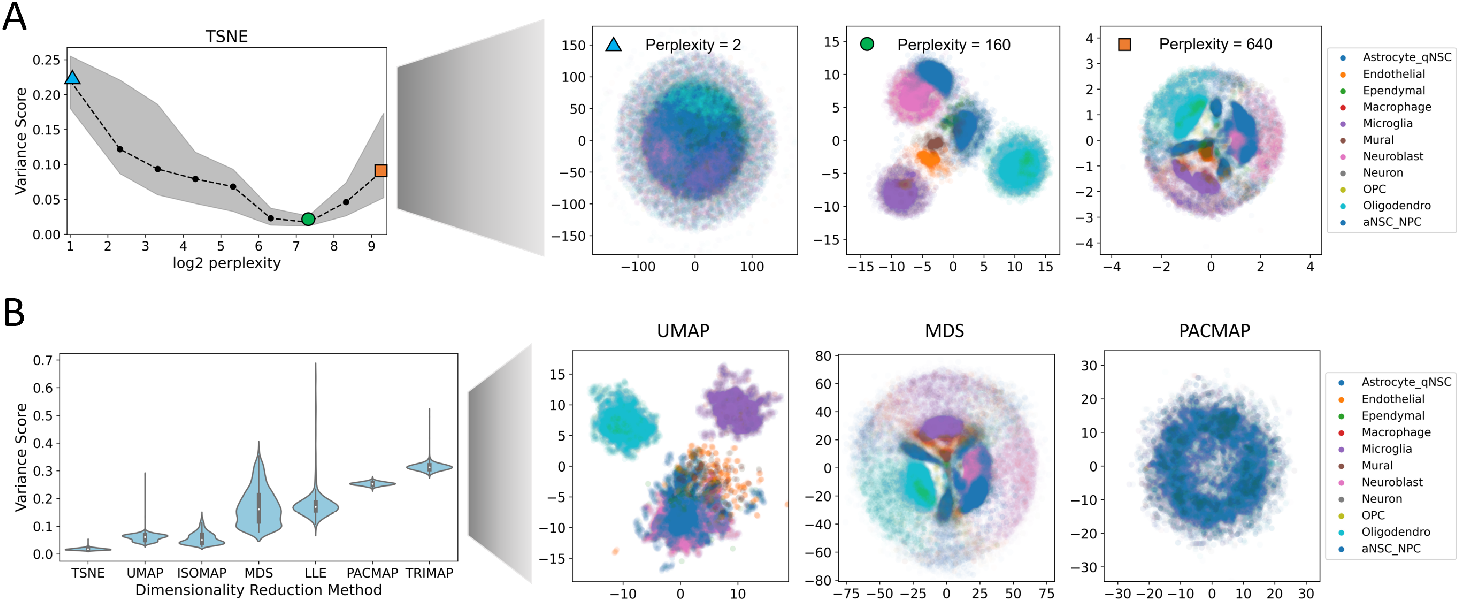
Optimizing DR algorithms using variance score. (A) Variance scores of t-SNE visualizations computed for different choices of perplexity with stacked visualizations of all bootstrap t-SNE visualizations for the optimal perplexity value (perplexity = 160) and two representative non-optimal cases (perplexity=2, perplexity=640). (B) Distribution of variance scores for the optimal hyperparameter choices (lowest variance score) of different DR algorithms (perplexity: t-SNE; number of neighbors: UMAP, ISOMAP, LLE, PACMAP; number of inliers: TRIMAP) shown with stacked visualizations of all bootstrap visualizations from three representative DR algorithms. Variance scores and dynamic visualizations were computed for 100 bootstrap samples.

For most data visualization tasks, there is often no clear choice of DR algorithm and many different methods are used in practice. For example, in single-cell biological data analysis, t-SNE and UMAP are the most commonly employed methods, but recently introduced DR algorithms like PaCMAP [4] and TriMap [3] among others [5, 14] may have advantages over t-SNE/UMAP in certain contexts. As such, users of DR visualization methods are often faced with the challenge of choosing one of many DR algorithms for data visualization. Again, we propose the variance score as a natural metric for selecting the DR algorithm that produces the most stable and robust visualizations. For the mouse SVZ single-cell transcriptomic data, t-SNE is the optimal DR algorithm with the lowest mean variance score in comparison to other optimally configured DR algorithms including UMAP, Isomap, MDS, LLE, PaCMAP, and TriMap (Figure 3B). Notably, stacked visualization of the bootstrap visualizations used in computing the variance scores shows better visual separation and stability of clusters for DR algorithms with lower variance scores (Figure 3B). PaCMAP and TriMap had the highest variance scores, which may be due to susceptibility to duplicate observations from bootstrap sampling and intrinsic randomness in the DR algorithm respectively [3, 4].

### 2.9 Dynamic visualization for RNA velocity trajectories

The study of cellular dynamics and subsequent prediction of cell fate trajectories is a topic of growing interest in single-cell transcriptomic data analysis [37]. Many such efforts have used the proportion of spliced to unspliced mRNA transcripts per gene to compute RNA velocity as a measure of cellular dynamics [18, 19]. However, RNA velocity estimates are typically noisy and common downstream analyses such as visualization or construction of velocity graphs often rely on low-dimensional projections of the high-dimensional RNA velocity vectors [18, 19, 38] and thus are subject to distortions. The application of the bootstrap approach underlying dynamic visualization and calculation of variance scores may bolster confidence in the interpretation of RNA velocity analyses.

RNA velocity analyses are especially useful for single-cell transcriptomic data of cells with multiple differentiation states or developmental pathways. A classic example is the segregation of progenitor cells into different endocrine subtypes during development of the pancreas in mammals [39], which is characterized by four terminal states denoted alpha, beta, gamma, and epsilon [19, 38]. Dynamic visualization of mouse single-cell transcriptomes from this lineage reveal significant variability in the global positioning of the gamma and epsilon terminal states (Figure 4A), which is corroborated by higher variance scores in these terminal states as compared to other states (Figure 4B). Dynamic visualization of the endocrine lineage and bootstrap construction of the UMAP-based velocity graph also permits the estimate of confidence intervals for velocity pseudotime values by computing multiple pseudotime values for the same cell across the set of bootstrap data (Figure 4C). Using this approach, we observe that uncertainty in pseudotime generally decreases further along the endocrine differentiation pathway, suggesting that pseudotime predictions may be more reliable for cells that have stronger commitment to differentiation (Figure 4D).

**Fig. 4.**
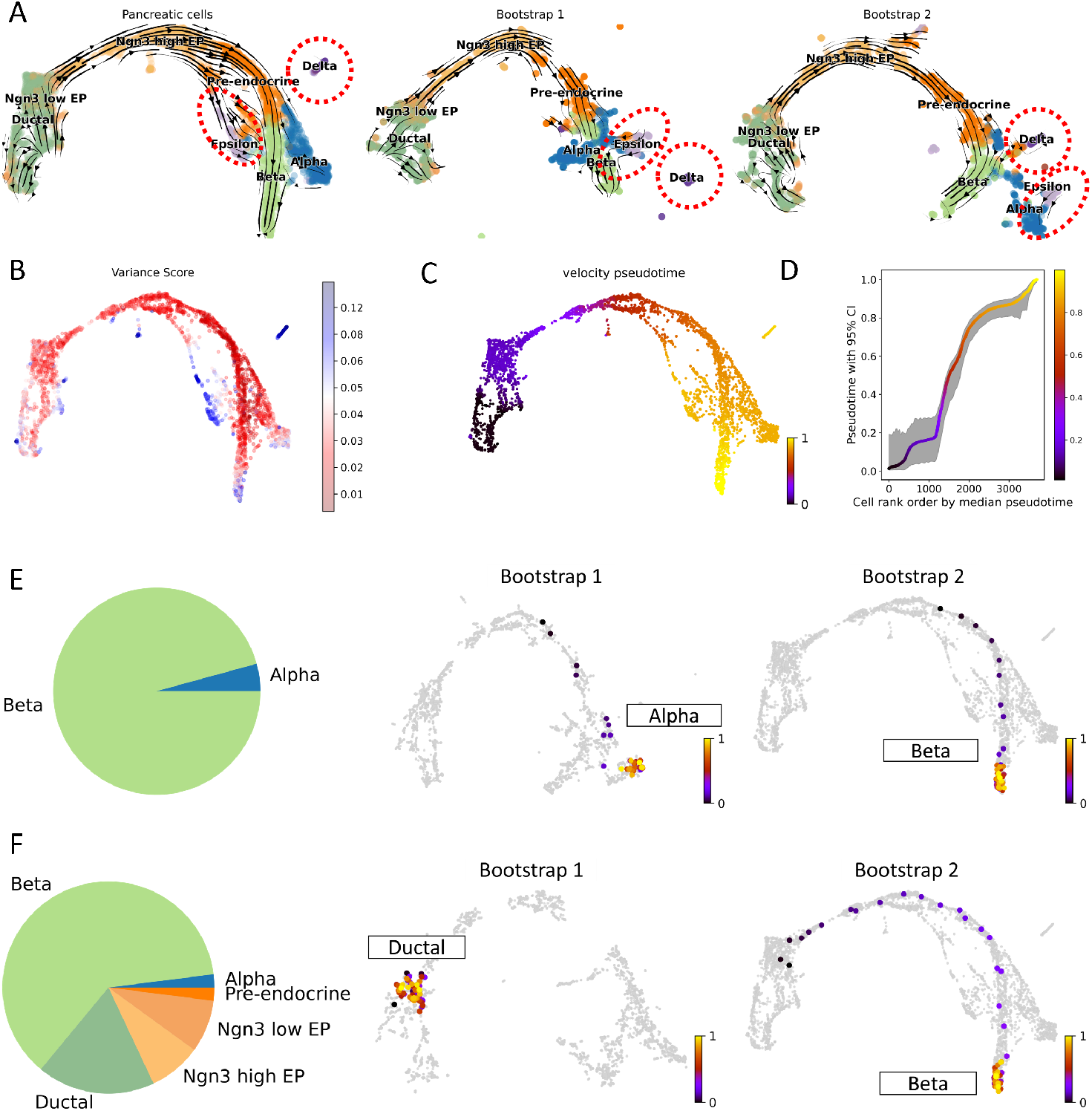
Application of dynamic visualization and variance score to RNA velocity analysis of single-cell transcriptomic data of mouse pancreas. (A) UMAP plots for the original data and two bootstrapped versions (left to right) shown with their associated RNA velocity embedding streams. Circled in red are the epsilon and delta cell clusters. (B) UMAP visualization of the original data with colors corresponding to variance score. Most high-variance cells are in the epsilon and delta clusters. (C) UMAP visualization of the original data with colors corresponding to RNA velocity pseudotime. (D) Rank-ordered pseudotimes computed for each cell over bootstrap UMAP visualizations with gray shading corresponding to 95% confidence interval. (E) Predicted terminal states of a Ngrn3 high EP cell using bootstrap UMAP-based velocity graphs and transitions traced across representative bootstrap UMAP visualizations for two terminal states (beta, alpha). Color corresponds to pseudotime along trajectory. (F) Trajectory analysis as in panel E except for a Ductal cell and with bootstrap visualizations for the two most common terminal states (beta, ductal) shown. Variance scores, pseudotimes, and trajectory predictions were computed for 100 bootstrap samples.

Given an initial cell, RNA velocity is often used to predict the possible terminal states of the cell. In many cases, these predictions are limited to a noisy walk across the velocity graph with only one terminal state identified. We leverage the same bootstrap approach as used in dynamic visualization to generate multiple UMAP-based velocity graphs, each with a terminal state prediction for the same cell. By combining these terminal state predictions with dynamic visualization, we present a comprehensive breakdown of possible terminal states and their likelihoods, represented by the fraction of bootstrap trajectories ending at that state. This approach can be used for different initial states. For example, a given Ngn 3 ^high^ cell is predicted to mostly differentiate into the beta terminal state (Figure 4E); a given ductal cell is predicted to mostly differentiate into the beta terminal state but has predictions targeting many other possible terminal states (Figure 4F). These applications are not limited to the endocrine lineage but are broadly applicable to other singlecell lineage data such as gastrulation of the erythroid lineage (Extended Data Figure A4) and human bone marrow development (Extended Data Figure A5).

## 3 Discussion

Our dynamic visualization procedure is compatible with any standard DR algorithm and can aid in the diagnosis of common interpretative pitfalls from viewing static DR visualizations. Importantly, the dynamic visualization procedure is as easy to understand and interpret as the underlying static DR algorithm, making it accessible to both technical and non-technical audiences. Dynamic visualization and variance score calculation are useful for improving confidence in interpretations of DR data visualization across a broad range of disciplines including but not limited to single-cell biology, spatial biology, genomics, and astronomy. As we have shown, dynamic visualizations can diagnose issues such as the stability of bridging connections between clusters, robustness of detected clusters, and order or patterns in labeled data visualizations–all of which can be impossible to diagnose under the current paradigm of static DR visualization. The variance score serves as a quality metric for DR visualizations that is complementary to existing concordance measures. The variance score has several desirable properties including positive correlation with variability among replicates of data and convergence to the variance score under real sampling of the data distribution for sufficiently large *n* or *B* and under regularity conditions.

Dynamic visualizations and variance scores are useful in many settings, which include optimizing hyperparameters, selecting the optimal DR algorithm, and assessing confidence in RNA velocity trajectory inference and pseudotime estimation.

Several metrics have been developed to measure the quality of DR visualizations with respect to preserving relationships in the original high-dimensional data. Scalar metrics quantify the quality of a visualization with a single summary statistic and include Kullback-Leibler divergence [40], trustworthiness [24], continuity [24], neighborhood hit [23], and normalized stress [41]. Unlike scalar metrics, point-wise metrics, which report a score for each observation in the visualization, can capture heterogeneities in quality within the same visualization [11]. These metrics largely fall under two classes. First, local quality scores measure the degree of preservation of an observation’s local neighborhood, which include Jaccard score [42], projection precision score [25], and ratios of pairwise distances in a neighborhood. Second, global quality scores measure the degree of preservation of an observation’s relationships with all other observations in the data, which include correlation between pairwise distance vectors [43], mean projection error [43], stretch [26], and compression [26]. EMBEDR, a recently introduced point-wise metric, provides both an quality metric and statistical significance estimates [14]. None of these methods measure the sensitivity of visualizations to perturbations in the data and/or stochasticity in the DR algorithm.

Despite the breadth of different metrics available for evaluating quality of DR visualizations, there is limited adoption of these metrics in research workflows and publications. We believe that this lack of usage and awareness is primarily due to two reasons. First, current quality scores are difficult to interpret, especially for users from non-technical backgrounds. Second, although current quality scores measure errors introduced by the DR method, they fail to capture variability and noise in the original high-dimensional data, which can be equally responsible for visualizations that are distorted from the truth. As such, the variance score is the first DR metric that accounts for both noise in the original data and for the effect of perturbations on distortions in the DR visualization. The variance score is complementary to existing quality metrics, which focus primarily on measuring the bias rather than the variance of the data generation and visualization process.

The current dynamic visualization procedure has several outstanding limitations. First, it is only directly applicable to DR algorithms that can handle input data with duplicate observations, which results from the bootstrap sampling approach. To extend dynamic visualizations to other methods, we have included some extensions in DynamicViz that include the option of adding low levels of noise to bootstrap sampled observations and performing subsampling of the original data in lieu of bootstrap sampling before DR visualization. Although these approaches can generate qualitatively similar results as compared to the bootstrap sample, they are implemented at the cost of increased complexity (e.g. size of subsamples, additional distortions from added noise). Designing optimal solutions for handling duplicate observations in DR visualizations may be a fruitful direction for future research. Similarly, both the dynamic visualization procedure and the calculation of variance scores requires generating a large number of DR visualizations, which scales linearly with the number of bootstrap samples and can be computationally expensive for visualizing data with many observations. To reduce computational overhead, the generation of bootstrap visualizations can be performed in parallel in DynamicViz. Finally, it should be noted that the theoretical guarantees of the bootstrap visualization are only valid under certain regularity conditions and these remain to be proven for individual DR algorithms. Addressing these limitations in the current dynamic visualization procedure will make both dynamic visualizations and variance scores applicable to a broader range of DR algorithms, data modalities, and research questions.

## 4 Methods

Here we present expanded explanations of the computational methods, theoretical considerations, additional utilities in DynamicViz, and details on data processing. The discussion of computational methods focuses firstly on the procedure for producing dynamic visualizations and secondly on the procedure for computing the variance score. We include further details on DR quality assessment metrics that use relationships between observations in the original high-dimensional data. We refer to these metrics as concordance scores and include several modified versions in DynamicViz.

### 4.1 Dimensionality reduction visualization methods

We used the scikit-learn (version 1.0.2) implementations of t-SNE, PCA, MDS, LLE, and Isomap. We used the umap-learn (version 0.5.2) implementation of UMAP. We used the trimap (version 1.1.2) implementation of TriMap and the pacmap (version 0.5.4) implementation of PaCMAP. Unless otherwise specified, we used the default hyperparameter values for all DR visualization methods with reduction to two dimensions.

### 4.2 Dynamic visualizations

The procedure for generating dynamic visualizations is illustrated in Figure 1A, and is succinctly summarized as the following steps:

1. Bootstrap sampling is used to generate bootstrap data from the original data *X*.
2. Visualizations are computed for each bootstrap data according to a chosen DR method.
3. Bootstrap visualizations are aligned to the original visualization using a rigid algorithm that preserves Euclidean distances between observations in the visualization.
4. The resulting set of aligned bootstrap visualizations are integrated into one of three viewing modalities: interactive, animated, and stacked.

#### 4.2.1 Constructing bootstrap visualizations

The first step in constructing bootstrap visualizations is to perform the standard DR visualization of the data *X* ∈ ℝ^*n*×*p*^, which yields a visualization *f* (*X*) = *Y* ∈ ℝ^*n*×2^. Next, we perform bootstrap sampling of *X* by sampling rows of *X* (observations) with replacement until a set of *n* rows are sampled. This sampling procedure is repeated *B* times to create a set of *B* bootstrap data {*X*^(1)^, …, *X*^(*B*)^}. Due to the nature of bootstrap sampling, there will likely be duplicate observations in the bootstrap data. Most implementations of DR methods are robust to duplicate observations or have built-in handling of duplicate observations. For those DR methods that have neither, we have included additional functionalities in the dynamic visualization implementation including subsampling of *n* − *m* out of *n* observations and the option of adding Gaussian noise to the bootstrap samples. The same bootstrap sampling is also applied to any metadata corresponding to the observations and to indices that map a corresponding bootstrap observation to its original row/index in *X*. For the given DR method *f* (·), visualizations are generated for the bootstrap samples, yielding the set {*f* (*X*^(1)^), …, *f* (*X*^(*B*)^)} = {*Y* ^(1)^, …, *Y* ^(*B*)^ }. Each of the bootstrap visualizations have the same dimensions as the original visualization *Y*.

In the code, the entire bootstrap visualization generation process is packaged into the `dynamicviz.boot.generate()` method, which takes the original data *X* and a given DR method or a string specifier for any of the built-in DynamicViz DR methods. Currently, implementations of UMAP, t-SNE, PCA, multi-dimensional scaling (MDS), locally linear embedding (LLE), and Isomap are available directly through DynamicViz. Users can specify the number of bootstrap samples, whether to compute a number of top principal components from PCA as an intermediate processing step, how much Gaussian noise to add to each dimension in a bootstrap sampled observation (default is none), and whether to fix random states for generating the bootstrap samples. For dynamic visualization purposes, we recommend more than 10 bootstrap samples and for variance score calculation, we recommend more than 50 bootstrap samples. Users can also specify the number of jobs to initialize for the bootstrap visualization procedure, which can decrease computation time.

#### 4.2.2 Rigid alignment of bootstrap visualizations

Many DR methods including t-SNE and UMAP minimize an objective function that is defined over the Euclidean distance (or *L*^2^ norm) between pairs of observations in the visualization, often in comparison to corresponding distances in the high-dimensional data [1, 2]. As a result, a visualization that minimizes a DR method’s objective function may be translated, rotated, or reflected without any change in optimality. We refer to these distance-preserving transformations as rigid transformations. This invariance is problematic for dynamic integration of the raw bootstrap visualizations because the result is unconstrained with respect to all possible rigid transformations for each independent bootstrap visualization. We establish constraints with two post-processing steps for the bootstrap visualization *Y* ^(*k*)^, *k* ∈ {1, …, *B*} : first, translation of *Y* ^(*k*)^ such that the centroid of *Y* ^(*k*)^ is equal to the centroid of *Y* ; and second, rotation and reflection of *Y* ^(*k*)^ to minimize the mean squared distance between concordant observations of *Y* ^(*k*)^ and *Y*.

In the first step, we define the centroid of the original visualization as 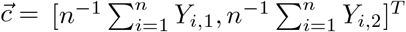 and the centroid of a bootstrap visualization as 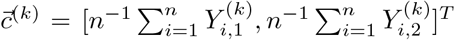. Then, the translated coordinates of an observation *i* in the bootstrap visualization is

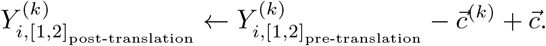

In practice, we translate both the original visualization and all bootstrap visualizations such that their centroids are equal to [0, 0] (i.e. shift the visualizations to the origin).

In the second step, we align the translated bootstrap visualization *Y* ^(*k*)^, *k* ∈ {1, …, *B*} obtained from the first step to the original visualization *Y* using the Kabsch algorithm, which seeks to minimize the mean squared distance between concordant observations of *Y* ^(*k*)^ and *Y* under three-dimensional rotation operations. The transformation used in the Kabsch algorithm is equivalent to rotations and reflections in two dimensions as long as the third dimension of the solution is equal to zero. In practice, we replace *Y* with 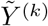, which is constructed by matching each observation in *Y* ^(*k*)^ with the embedding of the observation in *Y*. We use the `scipy.spatial.transform.Rotation.align_vectors()` implementation for the Kabsch algorithm to minimize the distance between *Z*^(*k*)^ and 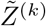, which are *Y* ^(*k*)^ and 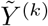 with zero-padded third axes. As such, the formulation of the Kabsch objective function can be represented as:

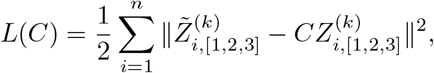

where we seek to minimize *L*(*C*) with respect to the three-dimensional rotation matrix *C*.

#### 4.2.3 Interactive visualization

Dynamic visualization with interactive integration of the bootstrap visualizations are implemented using the Plotly Dash framework and provide both inline interactive Jupyter displays and HTML versions of the visualizations that can be saved or embedded into web documents. In DynamicViz, interactive visualizations can be created directly from the output of `dynamicviz.boot.generate()` using the method `dynamicviz.viz.interactive()`. We use interpolation between bootstrap visualization frames and provide functionality for turning on/off the visibility of certain observations by their legend labels.

#### 4.2.4 Animated visualization

Dynamic visualization with animated integration of the bootstrap visualizations generates GIF (Graphics Interchange Format) or AVI (Audio Video Interweave) files. Each bootstrap visualization is presented as a frame in the file. Unlike interactive visualizations, there is no user interface for the final output of animated visualizations, but users can specify parameters related to the visualization and frame rate. This format is likely to be easier to embed in other media than interactive visualizations. In DynamicViz, interactive visualizations can be created directly from the output of `dynamicviz.boot.generate()` using the method `dynamicviz.viz.animated()`.

#### 4.2.5 Stacked visualization

Dynamic visualization with stacked integration of the bootstrap visualizations generates static PNG (Portable Network Graphics) files. Stacked visualization overlays all bootstrap visualizations with a user-specified opacity value and provides information that can be orthogonal to what is achievable using interactive or animated visualizations (e.g. for assessing robustness of patterns in labeled visualizations, see Figure 1FG). Stacked visualizations are particularly easy to incorporate into written scientific documents and presentations since image files are typically more flexible than HTML or video-based formats across most presentation modalities.

### 4.3 Variance score

Here, we outline some key properties and effects of different design choices on the variance score algorithm, which is defined according to Eq. (1) and reproduced below:

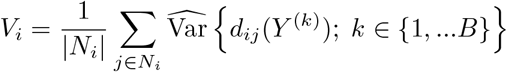

We focus on the use of different neighborhood structures in computing the variance score, different normalization schemes for the distances used in computing the variance score and their interpretations, transformations of the variance score that are available to users in DynamicViz, and the theoretical conditions for which the bootstrap is approximate to resampling from the underlying data distribution in the limit of sufficiently large *n* and *B*. The `dynamicviz.score.variance()` method is used to compute the variance score from the output of `dynamicviz.boot.generate()`.

#### 4.3.1 Neighborhood definition

The interpretation of the variance score of an observation indexed by *i* is sensitive to the definition of the neighborhood *N*_*i*_, which includes both global, local, and custom representations. In all examples shown thus far, we used the global neighborhood where an observation’s neighborhood is every other observation in the data. However, in practice, users of DynamicViz can use alternative definitions of neighborhoods to compute variance scores that are tailored to certain use cases. The neighborhood definition can be specified as an optional argument to `dynamicviz.score.variance()`.

For data with a large number of observations (i.e. large *n*), the global neighborhood variance score can be expensive to compute with complexity of *O*(*n*^2^). In these cases, we have implemented an alternative “random” computation of the global variance score, which randomly defines a neighborhood of size *k < n* for each observation such that the complexity reduces to *O*(*nk*). Importantly, the neighborhood definitions are not symmetric such that the observation at *j* being in the neighborhood of *N*_*i*_ does not imply that the observation at *I* is also in the neighborhood *N*_*j*_. Empirically, the “random” neighborhood definition can approximate the global neighborhood definition (Extended Data Figure A6).

When the local stability of the visualization embedding is more important than the global stability, the local neighborhood variance score can be used in lieu of the global version. An example of when the local variance score may be preferred over the global variance score is in visualizing highly clustered data with no assumption that the clusters are related to each other. The local neighborhood of *N*_*i*_ is defined as the *k* nearest observations to the observation at *i*, where the distance between observations is calculated as the Euclidean distance in the original high-dimensional data *X* and thus is consistently defined across all bootstrap visualizations. The local neighborhood definition is not symmetric.

For use cases where the global or local neighborhood definitions are insufficient, we have implemented an option to define symmetric neighborhoods (i.e. if observation at *j* is in *N*_*i*_, then observation at *i* is in *N*_*j*_). These symmetric neighborhoods are constructed from a list of discrete labels 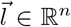 where observations with the same label are added to the same neighborhood.

The various neighborhood definitions outlined above can be summarized as follows:

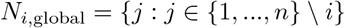

*N*_*i*,random_(*k*) = {*j* : *j* ∈ *R* containing *k* random elements drawn from {1, …, *n*} \ *i*}

*N*_*i*,local_(*k*) = {*j* : *j* ∈ *L* containing the *k* nearest neighbors of *i* in {1, …, *n*} \ *i*}

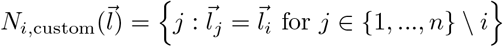

#### 4.3.2 Normalization of distances

Two primary normalization steps are used in computing the variance score. First, we optionally normalize the distance between an observation at *i* and its neighbor at *j* by the mean distance between the two observations across all *B* bootstrap visualizations. Performing this normalization step effectively yields a variance score that is independent of the mean distance between an observation and each of its neighbors, which is desirable in certain cases because the variance in distance increases in a quadratic fashion with the mean distance. Second, to ensure that the variance score is comparable across different DR methods and data, which generate visualizations at different scales, we normalize all distances by the mean distance across all observations and their neighborhoods. With this normalization, the variance score is comparable to the squared coefficient of variation in distance between observations and their neighbors. The default setting presented in our experiments and analyses is only using the empirical version of the second normalization:

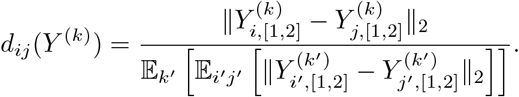

With the optional first normalization step included, the distance quantity becomes:

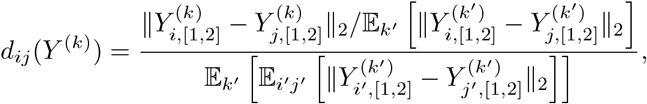

where 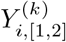 is two-dimensional vector describing the *i*-th observation of the visualization *Y* ^(*k*)^, 𝔼_*k*′_ is the expectation over the bootstrap visualizations indexed by *k*′, and 𝔼_*i*′ *j*′_ is the expectation over the observation indices *i*′ and *j*′.

#### 4.3.3 Transformation to stability score

In DynamicViz, we include a parametric transformation of the variance score, which we refer to as the stability score *S*. The stability score is computed as:

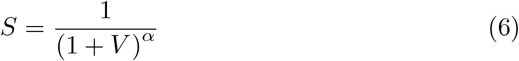

With the drawback of having to specify a parameter *α*, the stability score observes several desirable properties: first, the stability score spans a range between 0 and 1, which makes the stability score easy to interpret and translate across different settings; second, the stability score increases with more stable and robust embedding of observations, which yields the same general directionality as existing DR quality assessment metrics; third, usually for some values of *α*, symmetric distribution of stability scores can be achieved for a dataset. In practice, the stability score can be obtained by transforming the variance score using `dynamicviz.score.stability_from_variance()` or directly computed from the bootstrap visualizations output with `dynamicviz.score.stability()` for a value of *α*. We note that values of *α* between 5 and 50 are typically sufficient for achieving most of the desirable properties. The scaling of the color bar in Figure 4B was computed according to Eq. (6).

#### 4.3.4 Theoretical framework for the bootstrap and variance score

As is widely recognized in statistical theory, bootstrap samples can usually be regarded as surrogates of new samples generated from the same underlying distribution, which can be used to evaluate the uncertainty of a statistic of interest in an efficient and data-adaptive way.

Specifically, under mild regularity conditions on the visualization method *f* and the underlying distribution *P*_*X*_ of *X* (e.g., Section 3.2.2 of [44]), the empirical variance score defined in (2) for given (*i, j*) is a consistent estimator of the true conditional variance

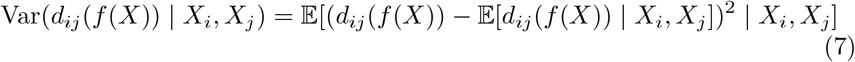

where the expectations on the right-hand side are taken over {*X*_1_, *X*_2_, …, *X*_*n*_} and the possibly random visualization function *f*, conditional on *X*_*i*_ and *X*_*j*_. Moreover, if the *n* samples in *X* are generated independently, the above conditional variance can also be written as

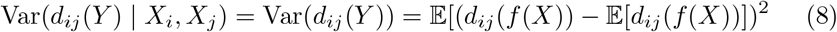

where on the right-hand side the expectation is with respect to the (*n* − 2) samples {*X*_1_, *X*_2_, …, *X*_*n*_}\{*X*_*i*_, *X*_*j*_}, and the visualization function *f*. In other words, the variance score estimates the variability of *d*_*ij*_(*f* (*X*)) if one applies *f* independently to replicated observations of {*X*_1_, …, *X*_*n*_} with only *X*_*i*_ and *X*_*j*_ fixed. These theoretical observations are also verified empirically in Figure 2BC and Extended Data Figure A1BC.

#### 4.3.5 Empirical validation of limiting properties of bootstrap

For both increasing *n* with fixed *B* and increasing *B* with fixed *n*, we show that the variance in Euclidean distance between pairs of observations across bootstrap visualizations approaches the variance in Euclidean distance of the same pairs of observations across visualizations of resampled data from the underlying data distribution (e.g. a mixture of Gaussian distributions; see Figure 2BC). We define the underlying data distribution as *P*_*X*_, from which we can sample observations *x* ∈ ℝ^*p*^. The procedure for generating visualizations for a given *n* and *B* is as follows:

1. 20 pairs of observations are drawn from *P*_*X*_
2. For each pair of observations defined (*i, j*):
  - For bootstrap visualizations, we draw *n* − 2 observations from *P*_*X*_ to construct *X*_context_ ∈ ℝ^(*n*−2)×*p*^. We bootstrap sample *X*_context_ for *B* times to get a set of bootstrap context data 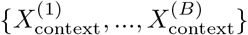.
  - For resampling visualizations, we repeatedly draw *n* − 2 observations from *G* for a total of *B* times to construct the set of context data 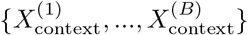.
  - We construct the set of data by concatenating the pair of observations to the context data: {*X*^(1)^, …, *X*^(*B*)^} where *X*^(*k*)^ ∈ ℝ^*n*×*p*^ for *k* ∈ {1, …, *B*}.
  - The set of data is used as input to a DR method *f* (·) to generate a set of visualizations {*Y* ^(*k*)^ : *Y* ^(*k*)^ = *f* (*X*^(*k*)^)} where *Y* ^(*k*)^ ∈ ℝ^*n*×2^. For each visualization, the resulting Euclidean distance between *i* and *j* in the visualization is recorded: ‖*Y*_*i*,[1,2]_ − *Y*_*j*,[1,2]‖2_.
  - Then, we compute 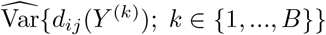 as described previously.
3. We compute the confidence intervals and mean of 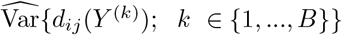 across all 20 pairs of observations.

### 4.4 Concordance scores

Existing approaches for quantifying the quality of a DR visualization are primarily concordannce scores, which measure the degree of agreement in local or global relationships between observations in the DR visualization and observations in the original high-dimensional data [11, 12]. Here, we present formulations of several concordance scores that are computed in Figure 2F to compare against the variance score. These formulations for the concordance scores are also available in DynamicViz.

#### 4.4.1 Spearman score

Spearman correlation *ρ* can be used to assess the agreement of rank-ordered distances between a given observation at *i* and all other observations in the visualization as compared to the original high-dimensional data [43]. The Spearman correlation between the Euclidean distances of an observation is computed using the `scipy.stats.spearmanr` implementation [45].

#### 4.4.2 Jaccard score

The Jaccard score or Jaccard set-distance for DR visualization assessment measures the degree of overlap between the *k*-nearest neighborhood of an observation in the visualization and its *k*-nearest neighborhood in the original high-dimensional data [42]. It is computed as the intersection cardinality over the union cardinality of the two neighborhoods:

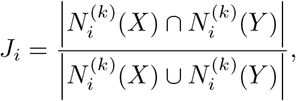

where 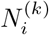 refers to the *k*-nearest neighborhood of observation at *i* using Euclidean distance.

#### 4.4.3 Mean projection error score

Mean projection error, also referred to as aggregate projection error, measures the average error of projection [43]. It is defined:

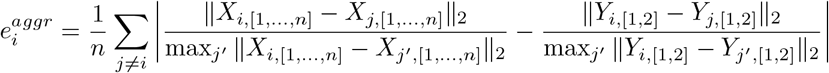

To define a concordance score (i.e. larger positive values indicating higher quality) based on the mean projection error that scales between 0 and 1, we compute:

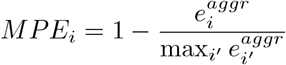

#### 4.4.4 Distortion score

We introduce a distortion score that is inspired by previous definitions of distortion in DR visualizations [13] and defined as the absolute value of the log fold change of the ratio between farthest-closest neighbors defined in the original high-dimensional data *X* and the ratio of the distances of the same neighbors in the visualization *Y* :

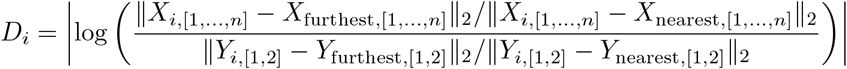

Importantly, the “farthest” and “closest” neighbors are defined exclusively on the original high-dimensional data *X*.

#### 4.4.5 Stretch score

Stretch measures the degree of stretching of pairwise distances in the DR visualization as compared to the original data [26], and is defined:

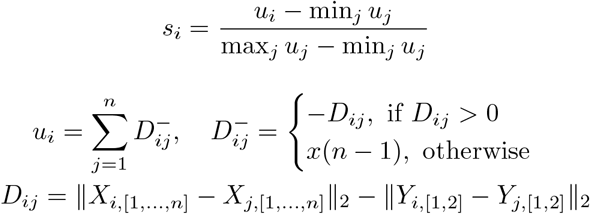

To define a concordance score (i.e. larger positive values indicating higher quality) based on stretch that scales between 0 and 1, we compute:

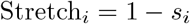

### 4.5 Data processing

#### 4.5.1 Synthetic data from mixture of Gaussian distributions

We are not aware of any large collections of real-world high-dimensional data that are drawn from the same underlying distribution. As such, to validate the theoretical guarantees of the bootstrap visualization and variance score, we generated synthetic data from a mixture of Gaussian distributions. The synthetic data permitted us to examine the empirical convergence of measures computed on bootstrap sampled data to the same measures computed on resampled data from the mixture of Gaussian distributions. We defined the mixture of Gaussian distributions as *k* multivariate Gaussian distributions with *p* dimensions. The mean of each Gaussian indexed *j* ∈ {1, …, *k*} was defined by a zero vector with the *j*-th element set to *m >* 0. The covariance matrix of each Gaussian was the identity matrix (i.e. no covariance between dimensions). We used values of *m* = 4, *k* = 5, and *p* = 50 for our experiments. An animated t-SNE visualization of this mixture of Gaussian distributions is available in Supplementary Video 5.

#### 4.5.2 Single-cell RNA-Seq, MERFISH, and CITE-Seq data processing

For single-cell RNA-seq data processing, we performed library size normalization by dividing the transcript count of each gene in a cell by the library size of that cell. A pseudocount was added to the normalized gene expression values and then we computed the logarithm of that value multiplied by 10^5^ as the final gene expression measure. Before DR visualization with t-SNE or UMAP, an initial DR was achieved by projecting the data to the top 50 principal components using PCA.

For the mouse subventricular zone data, we selected a random sample of 1,000 cells drawn from the single-cell RNA sequencing dataset of the subventricular zone, a neurogenic niche, from 28 male mice aged 3 months to 28 months old [27]. We used the 11 cell type labels identified in the original study [27]. The data is currently under embargo pending initial publication.

For the cell cycle data on mouse embryonic stem cells, we used a sample of 288 cells from the single-cell RNA sequencing dataset of mouse embryonic stem cells across three phases of cell cycle and 38,390 transcripts [32]. The data is publicly available at https://www.ebi.ac.uk/arrayexpress/experiments/E-MTAB-2805/.

For the human bone marrow data, we selected a random sample of 1,000 cells from the single-cell RNA sequencing dataset of cryopreserved bone marrow stem/progenitor CD34+ cells from healthy human donors [31]. We used the retrieved data from `scvelo.datasets.bonemarrow()`, which included 14,319 genes and 10 cell type labels (not all were represented in our sample). For the gastrulation data, we selected a random sample of 1,000 cells from the mouse embryo gastrulation single-cell RNA sequencing dataset restricted to the erythroid lineage [33]. We used the retrieved data from `scvelo.datasets.gastrulation_erythroid()`, which included 52,801 transcripts and 5 cell type labels.

For the pancreas data, which was exclusively used in the RNA velocity analyses, we used the entire dataset from single-cell RNA sequencing of mouse pancreatic epithelial and *NV F* ^+^ cells during secondary transition [39]. Unlike the previous data, the processing of the pancreas data used the default procedure outlined in the scvelo package [19]. The data was retrieved using `scvelo.datasets.pancreas()` and included 27998 genes across 3696 cells, spanning 8 cell types [39].

For the spatial transcriptomics data, we sampled 1,000 cells (or all available cells) for each of the 12 replicates in the MERFISH spatial transcriptomics dataset of mouse primary motor cortex [34]. The data was retrieved from the VizGen Resources web site at https://vizgen.com/resources/molecular-spatial-and-projection-diversity-of-neurons-in-primary-motor-cortex-revealed-by-in-situ-single-cell-transcriptomics/ and included 254 genes, spanning 24 cell types [34].

#### 4.5.3 Human genomics data from 1000 Genomes Project

We retrieved the genotype data from the 1000GP from http://ftp.1000genomes.ebi.ac.uk/vol1/ftp/release/20130502/supporting/hd_genotype_chip/ and used the same data processing as in [8]. We sampled 1,000 individuals from the genotype data that spanned 5 population clusters [28, 29].

#### 4.5.4 Astronomical object data from Sloan Digital Sky Survey

We used processed data from the Sloan Digital Sky Survey DR17 [30], which was retrieved from https://www.kaggle.com/datasets/fedesoriano/stellar-classification-dataset-sdss17. We sampled 2,000 observations at random and selected 8 relevant features related to the spectra measurements of each observation. The data spanned 3 classes of observations: star, quasar, and galaxy.

## Supplementary information

We include supplementary HTML files corresponding to the interactive visualization of Figure 1B (Supplementary Video1.html), Figure 1C (Supplementary Video1.html), and the mixture of Gaussian distributions (Supplementary Video5.html). We also include GIF files corresponding to animated visualization of Figure 1D (Supplementary Video3.gif) and Figure 1E (Supplementary Video4.gif).

## Acknowledgments

We would like to thank Kyle Swanson and Christine Yeh for their feedback on the functionality of DynamicViz. Fellowship support was provided by Knight-Hennessy Scholars program, Paul and Daisy Soros Fellowship for New Americans, and the National Science Foundation Graduate Research Fellowship Program to E.D.S. R.M. is supported by Professor David Donoho at Stanford University. J.Z. is supported by NSF CAREER 1942926, NIH P30AG059307, 5RM1HG010023 and grants from the Silicon Valley Foundation and the Chan-Zuckerberg Initiative.

## Data availability

The study primarily used publicly available data. Single-cell RNA-seq data on mouse subventricular zone is associated with a preprint and is currently under embargo pending initial publication. Single-cell RNA-seq data on mouse embryonic stem cells is available at https://www.ebi.ac.uk/arrayexpress/experiments/E-MTAB-2805/. Single-cell RNA-seq data human bone marrow is available through the Human Cell Atlas data portal: https://prod.data.humancellatlas.org/explore/projects/29f53b7e-071b-44b5-998a-0ae70d0229a4. Single-cell RNA-seq data on gastrulation of the erythroid lineage data can be downloaded according to instructions at https://github.com/MarioniLab/EmbryoTimecourse2018. Single-cell RNA-seq data on mouse pancreas lineage can be found on Gene Expression Omnibus under accession number GSE132188. The MERFISH spatial transcriptomics data on mouse primary motor cortex is available at the VizGen Resources web site at https://vizgen.com/resources/molecular-spatial-and-projection-diversity-of-neurons-in-primary-motor-cortex-revealed-by-in-situ-single-cell-transcriptomics/. The human bone marrow mononuclear cell CITE-Seq data is available through the Open Problems in Single-Cell Analysis Challenge at https://openproblems.bio/neurips_2021/. The genomic data from 1000 Genomes is available through http://ftp.1000genomes.ebi.ac.uk/vol1/ftp/release/20130502/supporting/hd_genotype_chip/. Processed data from the Sloan Digital Sky Survey DR17 was retrieved from https://www.kaggle.com/datasets/fedesoriano/stellar-classification-dataset-sdss17.

## Code availability

DynamicViz code is available at https://github.com/sunericd/dynamicviz.

## Author’s contributions

E.D.S. and J.Z. conceived of the study. E.D.S. designed and implemented the method with input from J.Z. and R.M. R.M. contributed to the theoretical framework for the study. E.D.S. prepared a draft of the manuscript. J.Z. and R.M. edited the manuscript.

## Extended Data A Supplemental Figures

**Fig. A1.**
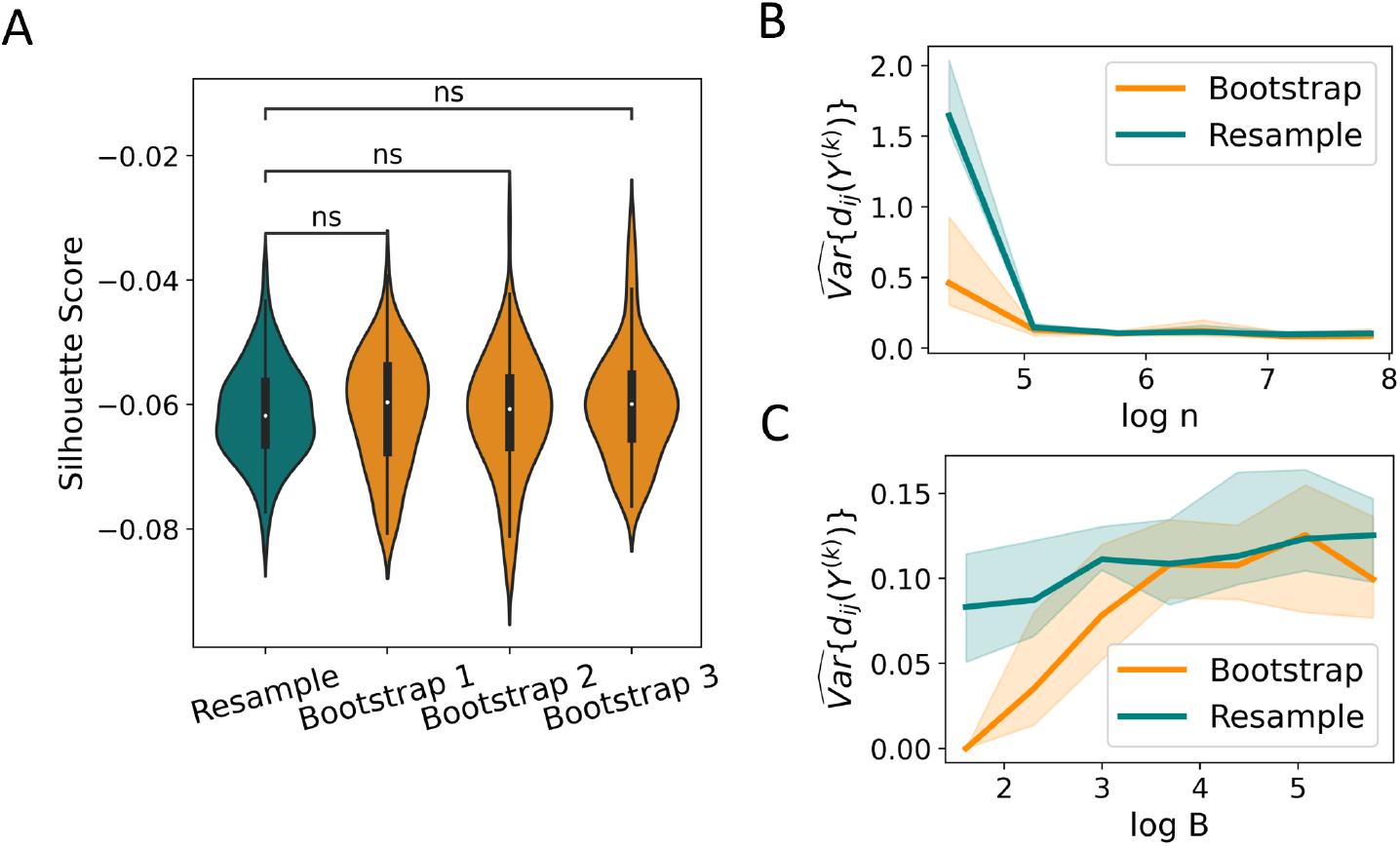
Empirical validation of bootstrap approximation to resampling in dynamic visualization. (A) Silhouette scores computed for t-SNE visualizations of data drawn repeatedly from a mixture of Gaussian distributions (100 repeated draws, *n*=500, *p*=100) in green compared to silhouette scores computed for t-SNE visualizations of 100 bootstrap samples from one initial sample from the mixture of Gaussian distributions in orange. Shown are bootstrap results for three representative initial samples. No statistically significant differences between any groups at Wilcoxon rank-sum test p-value cutoff of 0.05. (B) Variance in the Euclidean distance between two predetermined observations in t-SNE visualization across either 100 bootstrap sample data or 100 resampled data from a Gaussian mixture model (50 features, 5 distributions) for different numbers of observations, *n*. 95% confidence intervals are shown for 20 pairs of predetermined observations. (C) Same setting as panel C except with *n* = 1000 and different number of bootstrap samples or resamples, *B*. In accordance with bootstrap theory, the variance of pairwise distances across bootstrap visualizations approximates the variance of pairwise distances across resampled data from the same distribution as *n* increases and as *B* increases. Likewise, bootstrap visualizations produce qualitatively similar clustering patterns as visualization of resampled data from the same distribution, suggesting that both dynamic visualization and variance score can capture natural variability in data generation.

**Fig. A2.**
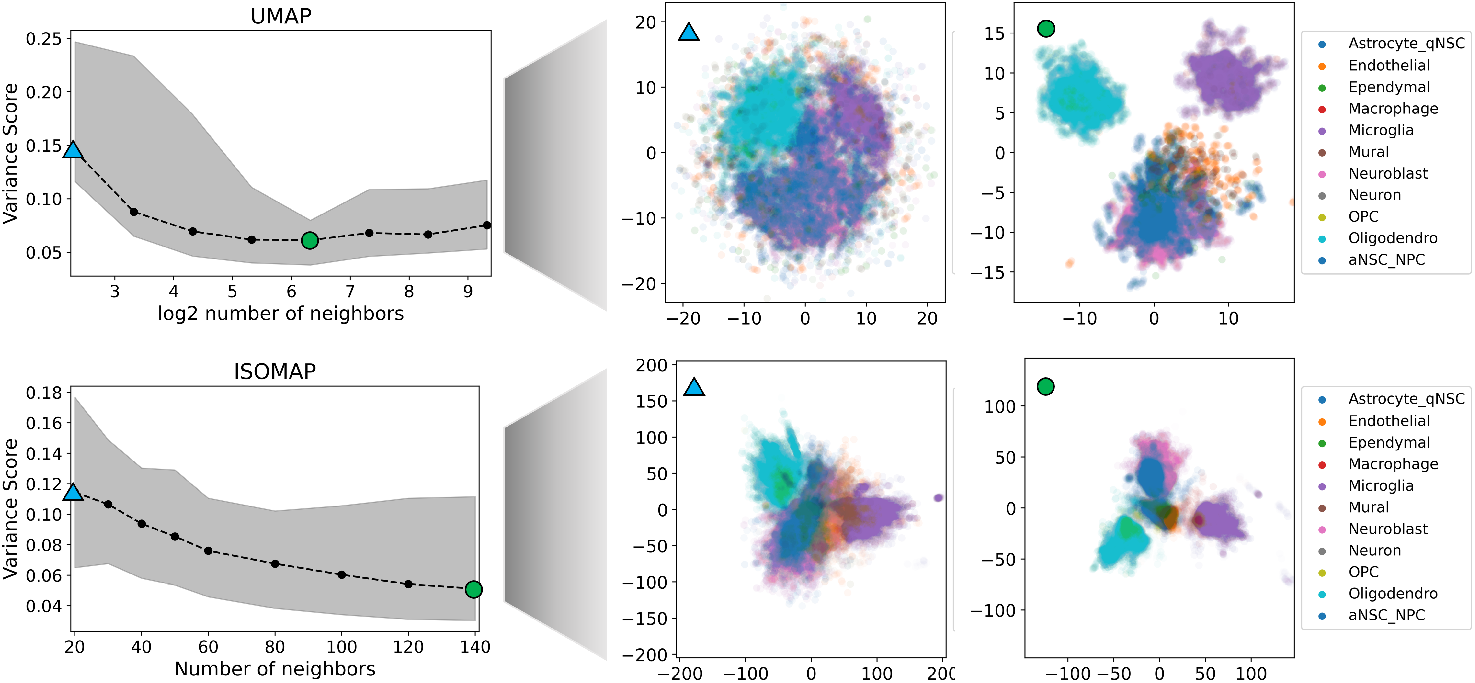
Variance scores of DR visualizations computed for different hyperparameter choices with stacked visualizations for the value with lowest variance score (green circle) and highest variance score (blue triangle). Shown are results for 100 bootstrap visualizations of the mouse subventricular zone single-cell transcriptomics data for (A) UMAP and (B) Isomap. Visualizations are not shown for LLE, PaCMAP, or TriMap since the variance scores were significantly higher for those methods. Generally, the hyperparameter values corresponding to minimal variance score also produced the most visually separated and stable cell type clusters.

**Fig. A3.**
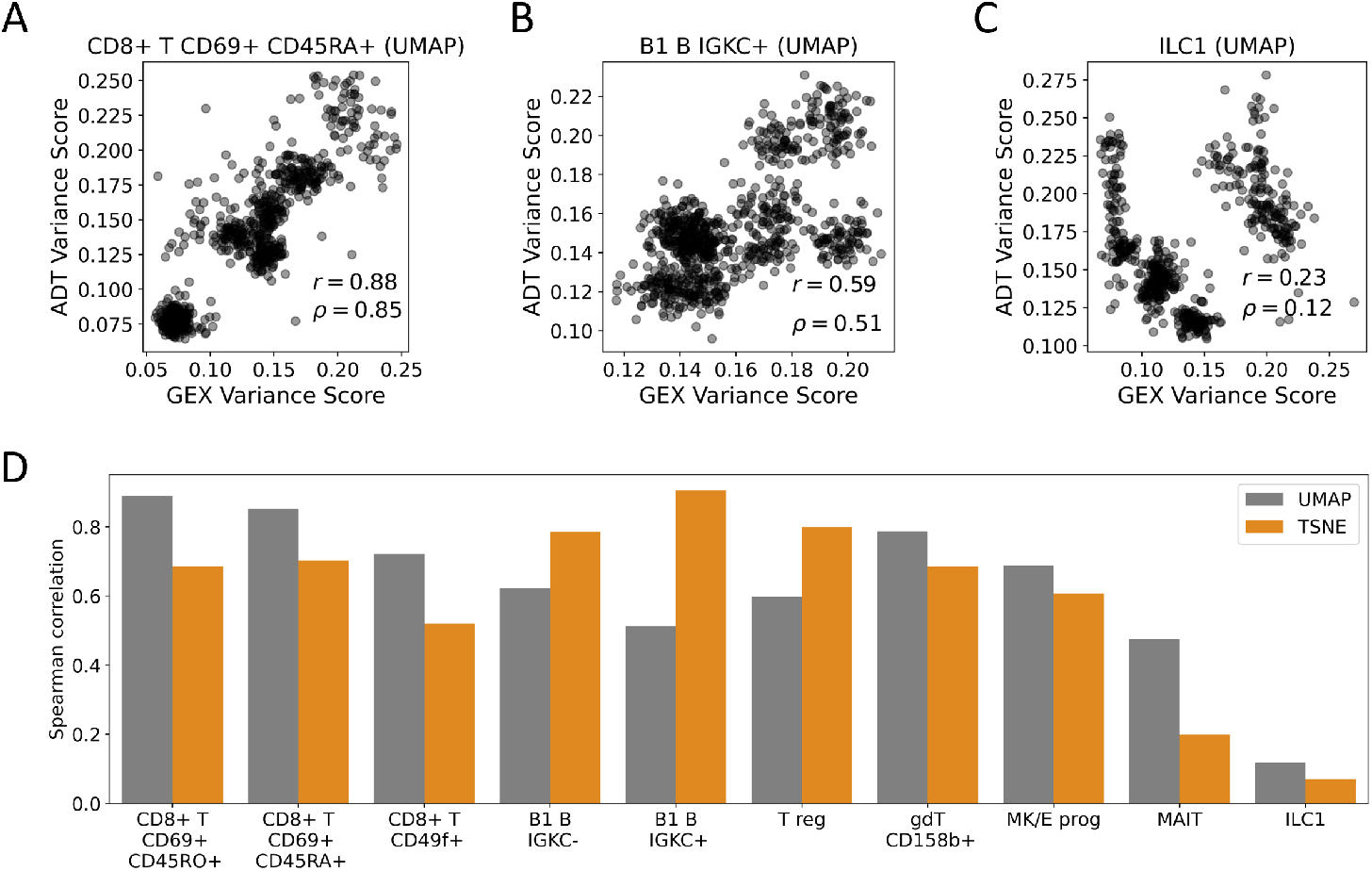
Correlation of variance scores between paired single-cell surface proteomic and transcriptomic signatures in CITE-seq data of human bone marrow. (A) Variance scores computed from UMAP visualization of CITE-seq transcriptomic data and variance scores computed from UMAP visualization of CITE-seq surface proteomic data for CD8+ T CD69+ CD45RA+ cells; (B) for B1 B IGKC+ cells; (C) and for ILC1 cells. (D) Spearman correlation between variance scores computed from DR visualizations of CITE-seq transcriptomic data and variance scores computed from DR visualizations of CITE-seq surface proteomic data for 10 human bone marrow cell types. See Appendix B for a discussion of these results.

**Fig. A4.**
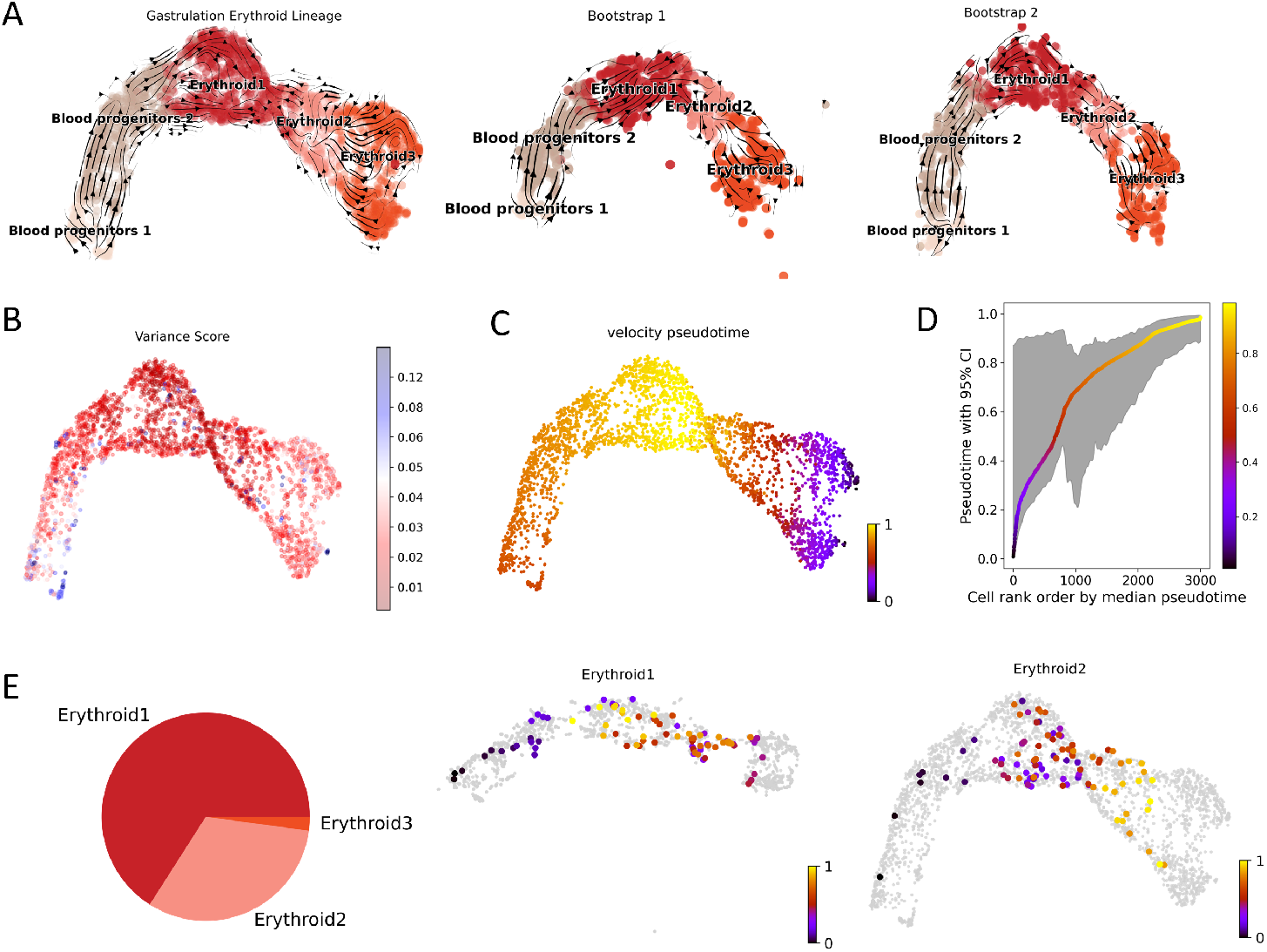
Application of dynamic visualizations and variance scores to RNA velocity analysis of single-cell data for gastrulation of the erythroid lineage. (A) RNA velocity embedding stream UMAP plots for the original data and two bootstrapped versions (left to right). UMAP visualization of the original data with colors corresponding to variance score. UMAP visualization of the original data with colors corresponding to RNA velocity pseudotime. (D) Rank-ordered pseudotimes computed for each cell over bootstrap UMAP visualizations with gray shading corresponding to 95% confidence interval. (E) Predicted terminal states of a Blood Progenitor 1 cell using RNA velocity trajectory analysis across bootstrap UMAP visualizations and transitions traced across representative bootstrap visualizations for two terminal states (Erythroid1. Erythroid2). Color corresponds to pseudotime along trajectory. Dynamic visualization and variance score provide a more detailed picture of standard RNA velocity analyses in the gastrulation of erythroid lineage, including the stability of RNA velocity streams, cell fates, and pseudotime.

**Fig. A5.**
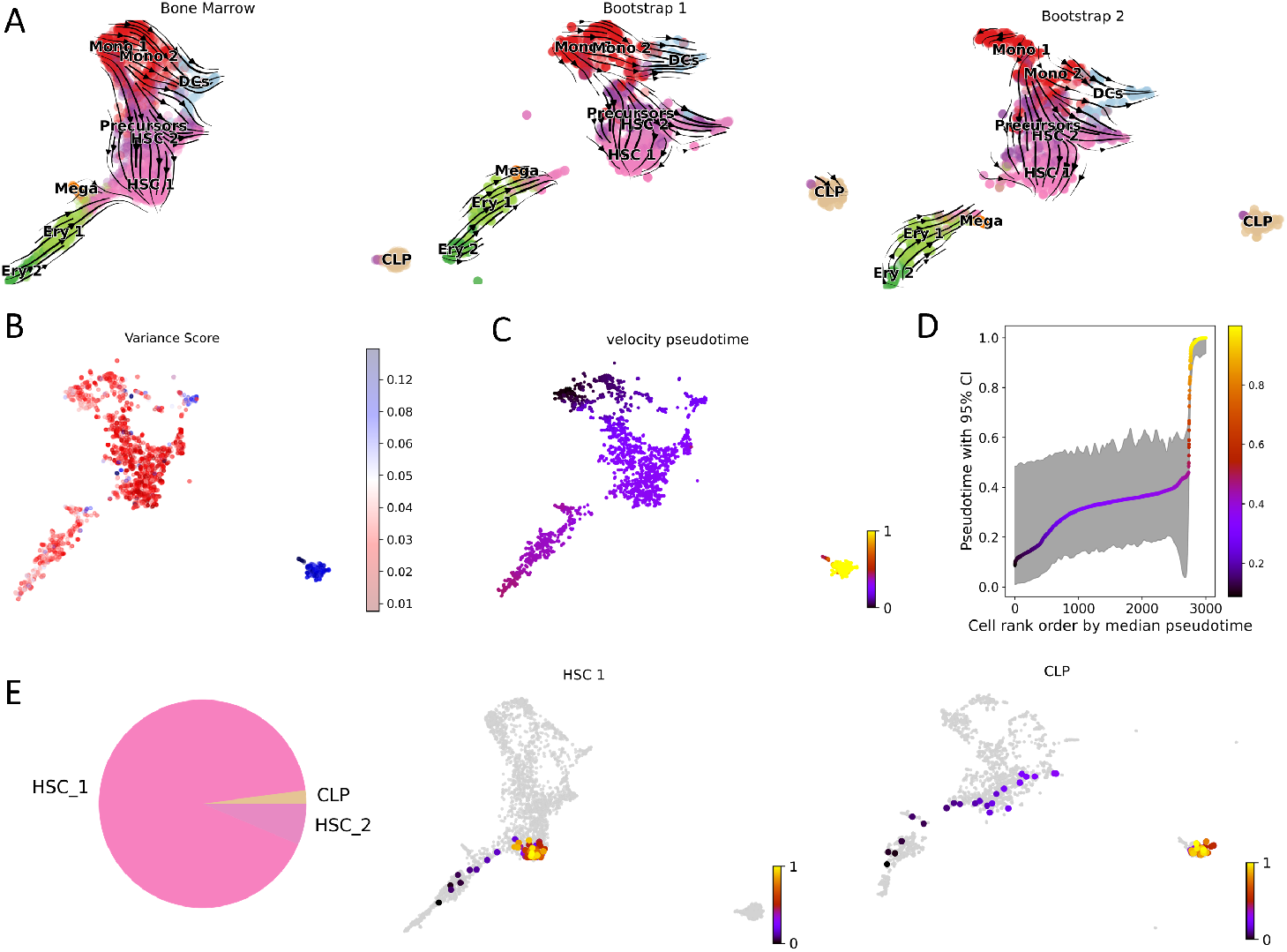
Application of dynamic visualizations and variance scores to RNA velocity analysis of single-cell data of human bone marrow. (A) RNA velocity embedding stream UMAP plots for the original data and two bootstrapped versions (left to right). (B) UMAP visualization of the original data with colors corresponding to variance score. (C) UMAP visualization of the original data with colors corresponding to RNA velocity pseudotime. (D) Rank-ordered pseudotimes computed for each cell over bootstrap UMAP visualizations with gray shading corresponding to 95% confidence interval. (E) Predicted terminal states of a Ery2 cell using RNA velocity trajectory analysis across bootstrap UMAP visualizations and transitions traced across representative bootstrap visualizations for two terminal states (HSC 1, CLP). Color corresponds to pseudotime along trajectory. Dynamic visualization and variance score provide a more detailed picture of standard RNA velocity analyses in the human bone marrow lineage, including the stability of RNA velocity streams, cell fates, and pseudotime.

**Fig. A6.**
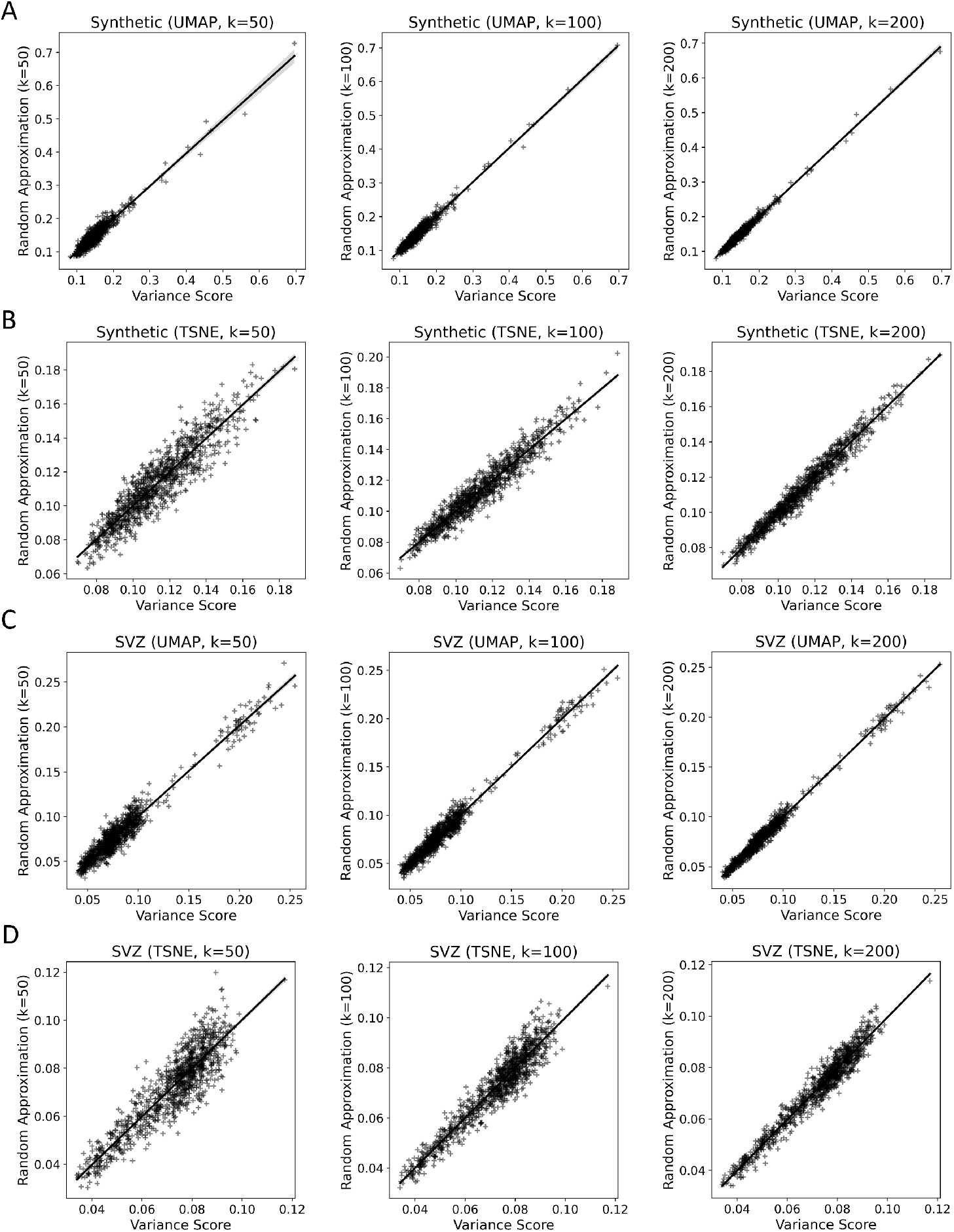
Random approximation of the global variance score for different choices of the number of random neighbors used, *k*, and for different combinations of DR visualization methods and data. (A) UMAP visualizations with synthetic data drawn from a mixture of Gaussians (n=1000, p=50). (B) t-SNE visualizations for the same synthetic data. (C) UMAP visualizations for the mouse subventricular zone (SVZ) single-cell transcriptomics data. (D) t-SNE visualizations for the same SVZ data. Generally, the random method for computing variance scores (see Methods for details) is a good approximation of the global variance score and the quality of this approximation increases with *k*.

## Extended Data B Variance score is correlated across data modalities

A key question is whether the variance score of an observation is correlated for visualizations on different sets of features. We investigate this property in the case of integrated transcriptomic and proteomic signatures in human bone marrow cells obtained using CITE-seq [46], which performs single-cell RNA sequencing in tandem with measurement of cell surface proteins [47]. For both t-SNE and UMAP dynamic visualizations, the variance scores of the transcriptomic data was positively correlated with the variance scores for surface proteomic data across all 10 cell types included in this analysis (Extended Data Figure A3), suggesting that variance scores obtained from either transcriptomic or proteomic data of a cell may capture variability in the DR visualization of a shared underlying biological signal. Generally, we found that T cells and B cells, which are composed of several heterogeneous subtypes, tend to have higher conservation of variance scores between transcriptomic and surface proteomic visualizations than other cell types. These results indicate that the variance score may quantify the intrinsic stability of clustered data visualizations irrespective of the features used to generate the visualizations. For the CITE-seq data with paired transcriptomic and surface proteomic measurements, we used the subset of 10 cell types for which there were between 500 and 1,000 total cells per cell type in the CITE-Seq dataset from human bone marrow mononuclear cells [46]. Each cell had expression values for 13,953 genes and 134 surface proteins. The data is publicly available through the Open Problems in Single-Cell Analysis Challenge at https://openproblems.bio/neurips_2021/.

## References

[1] van der Maaten, L. J. P. & Hinton, G. E. Visualizing High-Dimensional Data Using t-SNE. Journal of Machine Learning Research 9 (nov), 2579–2605 (2008). Publisher: Microtome Publishing.

[2] McInnes, L., Healy, J. & Melville, J. UMAP: Uniform Manifold Approximation and Projection for Dimension Reduction. 1802.03426 [cs, stat] (2020). URL http://arxiv.org/abs/1802.03426. ArXiv: 1802.03426.

[3] Amid, E. & Warmuth, M. K. TriMap: Large-scale Dimensionality Reduction Using Triplets. 1910.00204 [cs, stat] (2019). URL http://arxiv.org/abs/1910.00204. ArXiv: 1910.00204.

[4] Wang, Y., Huang, H., Rudin, C. & Shaposhnik, Y. Understanding How Dimension Reduction Tools Work: An Empirical Approach to Deciphering t-SNE, UMAP, TriMap, and PaCMAP for Data Visualization. Journal of Machine Learning Research 22 (201), 1–73 (2021). URL http://jmlr.org/papers/v22/20-1061.html.

[5] Moon, K. R. et al. Visualizing Structure and Transitions for Biological Data Exploration. preprint, Bioinformatics (2017). URL http://biorxiv.org/lookup/doi/10.1101/120378.

[6] Kobak, D. & Berens, P. The art of using t-SNE for single-cell transcriptomics. Nature Communications 10 (1), 5416 (2019). URL https://www.nature.com/articles/s41467-019-13056-x. https://doi.org/10.1038/s41467-019-13056-x, number: 1 Publisher: Nature Publishing Group.

[7] Su, Y., Shi, Q. & Wei, W. Single cell proteomics in biomedicine: High-dimensional data acquisition, visualization, and analysis. PROTEOMICS 17 (3-4), 1600267 (2017). URL https://onlinelibrary.wiley.com/doi/abs/10.1002/pmic.201600267. https://doi.org/10.1002/pmic.201600267, eprint: https://onlinelibrary.wiley.com/doi/pdf/10.1002/pmic.201600267.

[8] Diaz-Papkovich, A., Anderson-Trocme, L. & Gravel, S. A review of UMAP in population genetics. Journal of Human Genetics 66 (1), 85–91 (2021). URL https://www.nature.com/articles/s10038-020-00851-4. https://doi.org/10.1038/s10038-020-00851-4, number: 1 Publisher: Nature Publishing Group.

[9] Anders, F. et al. Dissecting stellar chemical abundance space with t-SNE. Astronomy & Astrophysics 619, A125 (2018). URL http://arxiv.org/abs/1803.09341. https://doi.org/10.1051/0004-6361/201833099, 1803.09341.

[10] Cooley, S. M., Hamilton, T., Aragones, S. D., Ray, J. C. J. & Deeds, E. J. A novel metric reveals previously unrecognized distortion in dimensionality reduction of scRNA-seq data. Tech. Rep., bioRxiv (2022). URL https://www.biorxiv.org/content/10.1101/689851v6. Section: New Results Type: article.

[11] Espadoto, M., Martins, R. M., Kerren, A., Hirata, N. S. T. & Telea, A. C. Toward a Quantitative Survey of Dimension Reduction Techniques. IEEE Transactions on Visualization and Computer Graphics 27 (3), 2153–2173 (2021). https://doi.org/10.1109/TVCG.2019.2944182, conference Name: IEEE Transactions on Visualization and Computer Graphics.

[12] Nonato, L. G. & Aupetit, M. Multidimensional Projection for Visual Analytics: Linking Techniques with Distortions, Tasks, and Layout Enrichment. IEEE Transactions on Visualization and Computer Graphics 25 (8), 2650–2673 (2019). https://doi.org/10.1109/TVCG.2018.2846735, conference Name: IEEE Transactions on Visualization and Computer Graphics.

[13] Chari, T., Banerjee, J. & Pachter, L. The Specious Art of Single-Cell Genomics. Tech. Rep., bioRxiv (2021). URL https://www.biorxiv.org/content/10.1101/2021.08.25.457696v3. Section: New Results Type: article.

[14] Johnson, E. M., Kath, W. & Mani, M. EMBEDR: Distinguishing signal from noise in single-cell omics data. Patterns 3 (3), 100443 (2022). URL https://www.sciencedirect.com/science/article/pii/S2666389922000162. https://doi.org/10.1016/j.patter.2022.100443.

[15] Hao, Y. et al. Integrated analysis of multimodal single-cell data. Cell 184 (13), 3573–3587.e29 (2021). URL https://www.sciencedirect.com/science/article/pii/S0092867421005833. https://doi.org/10.1016/j.cell.2021.04.048.

[16] Stuart, T. et al. Comprehensive Integration of Single-Cell Data. Cell 177 (7), 1888–1902.e21 (2019). URL https://www.sciencedirect.com/science/article/pii/S0092867419305598. https://doi.org/10.1016/j.cell.2019.05.031.

[17] Butler, A., Hoffman, P., Smibert, P., Papalexi, E. & Satija, R. Integrating single-cell transcriptomic data across different conditions, technologies, and species. Nature Biotechnology 36 (5), 411–420 (2018). URL https://www.nature.com/articles/nbt.4096. https://doi.org/10.1038/nbt.4096, number: 5 Publisher: Nature Publishing Group.

[18] La Manno, G. et al. RNA velocity of single cells. Nature 560 (7719), 494–498 (2018). URL https://www.nature.com/articles/s41586%E2%80%93018%E2%80%930414%E2%80%936. https://doi.org/10.1038/s41586-018-0414-6, number: 7719 Publisher: Nature Publishing Group.

[19] Bergen, V., Lange, M., Peidli, S., Wolf, F. A. & Theis, F. J. Generalizing RNA velocity to transient cell states through dynamical modeling. Nature Biotechnology 38 (12), 1408–1414 (2020). URL https://www.nature.com/articles/s41587-020-0591-3. https://doi.org/10.1038/s41587-020-0591-3, number: 12 Publisher: Nature Publishing Group.

[20] Wattenberg, M., Viégas, F. & Johnson, I. How to Use t-SNE Effectively. Distill 1 (10), e2 (2016). URL http://distill.pub/2016/misread-tsne. https://doi.org/10.23915/distill.00002.

[21] Cooley, S. M. Distortion in Dimensionality Reduction and Implications for the Analysis of Single Cell RNA-Sequencing Data. Ph.D., University of California, Los Angeles, United States – California (2021). URL https://www.proquest.com/docview/2571111018/abstract/1C4D093B947C4AC5PQ/1. ISBN: 9798538118793.

[22] Wu, Y., Tamayo, P. & Zhang, K. Visualizing and Interpreting Single-Cell Gene Expression Datasets with Similarity Weighted Nonnegative Embedding. Cell Systems 7 (6), 656–666.e4 (2018). URL https://www.sciencedirect.com/science/article/pii/S240547121830440X. https://doi.org/10.1016/j.cels.2018.10.015.

[23] Paulovich, F. V., Nonato, L. G., Minghim, R. & Levkowitz, H. Least Square Projection: A Fast High-Precision Multidimensional Projection Technique and Its Application to Document Mapping. IEEE Transactions on Visualization and Computer Graphics 14 (3), 564–575 (2008). https://doi.org/10.1109/TVCG.2007.70443, conference Name: IEEE Transactions on Visualization and Computer Graphics.

[24] Venna, J. & Kaski, S. Visualizing gene interaction graphs with local multidimensional scaling. Proceedings of ESANN’06, 14th European Symposium on Artificial Neural Networks 557–562 (2006). Publisher: d-side group.

[25] Schreck, T., von Landesberger, T. & Bremm, S. Techniques for Precision-Based Visual Analysis of Projected Data. Information Visualization 9 (3), 181–193 (2010). URL https://doi.org/10.1057/ivs.2010.2. https://doi.org/10.1057/ivs.2010.2, publisher: SAGE Publications.

[26] Aupetit, M. Visualizing distortions and recovering topology in continuous projection techniques. Neurocomputing 70 (7), 1304–1330 (2007). URL https://www.sciencedirect.com/science/article/pii/S0925231206004814. https://doi.org/10.1016/j.neucom.2006.11.018.

[27] Buckley, M. T. et al. Cell type-specific aging clocks to quantify aging and rejuvenation in regenerative regions of the brain. Tech. Rep., bioRxiv (2022). URL https://www.biorxiv.org/content/10.1101/2022.01.10.475747v2. Section: New Results Type: article.

[28] Auton, A. et al. A global reference for human genetic variation. Nature 526 (7571), 68–74 (2015). URL https://www.nature.com/articles/nature15393. https://doi.org/10.1038/nature15393, number: 7571 Publisher: Nature Publishing Group.

[29] McVean, G. A. et al. An integrated map of genetic variation from 1,092 human genomes. Nature 491 (7422), 56–65 (2012). URL https://www.nature.com/articles/nature11632. https://doi.org/10.1038/nature11632, number: 7422 Publisher: Nature Publishing Group.

[30] York, D. G. et al. The Sloan Digital Sky Survey: Technical Summary. The Astronomical Journal 120 (3), 1579–1587 (2000). URL https://doi.org/10.1086/301513. https://doi.org/10.1086/301513, publisher: American Astronomical Society.

[31] Setty, M. et al. Characterization of cell fate probabilities in singlecell data with Palantir. Nature Biotechnology 37 (4), 451–460 (2019). URL https://www.nature.com/articles/s41587-019-0068-4. https://doi.org/10.1038/s41587-019-0068-4, number: 4 Publisher: Nature Publishing Group.

[32] Buettner, F. et al. Computational analysis of cell-to-cell heterogeneity in single-cell RNA-sequencing data reveals hidden subpopulations of cells. Nature Biotechnology 33 (2), 155–160 (2015). URL https://www.nature.com/articles/nbt.3102. https://doi.org/10.1038/nbt.3102, number: 2 Publisher: Nature Publishing Group.

[33] Pijuan-Sala, B. et al. A single-cell molecular map of mouse gastrulation and early organogenesis. Nature 566 (7745), 490–495 (2019). URL https://www.nature.com/articles/s41586-019-0933-9. https://doi.org/10.1038/s41586-019-0933-9, number: 7745 Publisher: Nature Publishing Group.

[34] Zhang, M. et al. Spatially resolved cell atlas of the mouse primary motor cortex by MERFISH. Nature 598 (7879), 137–143 (2021). URL https://www.nature.com/articles/s41586-021-03705-x. https://doi.org/10.1038/s41586-021-03705-x, number: 7879 Publisher: Nature Publishing Group.

[35] Cao, Y. & Wang, L. Automatic Selection of t-SNE Perplexity. 1708.03229 [cs, stat] (2017). URL http://arxiv.org/abs/1708.03229. ArXiv: 1708.03229.

[36] Belkina, A. C. et al. Automated optimized parameters for T-distributed stochastic neighbor embedding improve visualization and analysis of large datasets. Nature Communications 10 (1), 5415 (2019). URL https://www.nature.com/articles/s41467-019-13055-y. https://doi.org/10.1038/s41467-019-13055-y, number: 1 Publisher: Nature Publishing Group.

[37] Bergen, V., Soldatov, R. A., Kharchenko, P. V. & Theis, F. J. RNA velocity-current challenges and future perspectives. Molecular Systems Biology 17 (8), e10282 (2021). https://doi.org/10.15252/msb.202110282.

[38] Lange, M. et al. CellRank for directed single-cell fate mapping. Nature Methods 19 (2), 159–170 (2022). URL https://www.nature.com/articles/s41592-021-01346-6. https://doi.org/10.1038/s41592-021-01346-6, number: 2 Publisher: Nature Publishing Group.

[39] Bastidas-Ponce, A. et al. Comprehensive single cell mRNA profiling reveals a detailed roadmap for pancreatic endocrinogenesis. Development 146 (12), dev173849 (2019). URL https://doi.org/10.1242/dev.173849. https://doi.org/10.1242/dev.173849.

[40] Hinton, G. E. & Roweis, S. Becker, S., Thrun, S. & Obermayer, K. (eds) Stochastic Neighbor Embedding. (eds Becker, S., Thrun, S. & Obermayer, K.) Advances in Neural Information Processing Systems, Vol. 15 (MIT Press, 2002). URL https://proceedings.neurips.cc/paper/2002/file/6150ccc6069bea6b5716254057a194ef-Paper.pdf.

[41] Joia, P., Coimbra, D., Cuminato, J. A., Paulovich, F. V. & Nonato, L. G. Local Affine Multidimensional Projection. IEEE Transactions on Visualization and Computer Graphics 17 (12), 2563–2571 (2011). https://doi.org/10.1109/TVCG.2011.220, conference Name: IEEE Transactions on Visualization and Computer Graphics.

[42] Martins, R. M., Minghim, R. & Telea, A. C. Borgo, R. & Turkay, C. (eds) Explaining Neighborhood Preservation for Multidimensional Projections. (eds Borgo, R. & Turkay, C.) Computer Graphics and Visual Computing (CGVC) (The Eurographics Association, 2015).

[43] Martins, R. M., Coimbra, D. B., Minghim, R. & Telea, A. C. Visual analysis of dimensionality reduction quality for parameterized projections. Computers & Graphics 41, 26–42 (2014). URL https://www.sciencedirect.com/science/article/pii/S0097849314000235. https://doi.org/10.1016/j.cag.2014.01.006.

[44] Shao, J. & Tu, D. The Jackknife and Bootstrap Springer Series in Statistics (Springer, New York, NY, 1995). URL http://link.springer.com/10.1007/978-1-4612-0795-5.

[45] Kokoska, S. & Zwillinger, D. CRC Standard Probability and Statistics Tables and Formulae, Student Edition (CRC Press, 2000). Google-Books-ID: G5hJqwjweiUC.

[46] Luecken, M. et al. Vanschoren, J. & Yeung, S. (eds) A sandbox for prediction and integration of DNA, RNA, and proteins in single cells. (eds Vanschoren, J. & Yeung, S.) Proceedings of the Neural Information Processing Systems Track on Datasets and Benchmarks, Vol. 1 (2021). URL https://datasets-benchmarks-proceedings.neurips.cc/paper/2021/file/158f3069a435b314a80bdcb024f8e422-Paper-round2.pdf.

[47] Stoeckius, M. et al. Simultaneous epitope and transcriptome measurement in single cells. Nature Methods 14 (9), 865–868 (2017). URL https://www.nature.com/articles/nmeth.4380. https://doi.org/10.1038/nmeth.4380, number: 9 Publisher: Nature Publishing Group.

