## Supplementary figures and images for "Dynamic visualization of high-dimensional data"

### Supplementary_Video3.gif

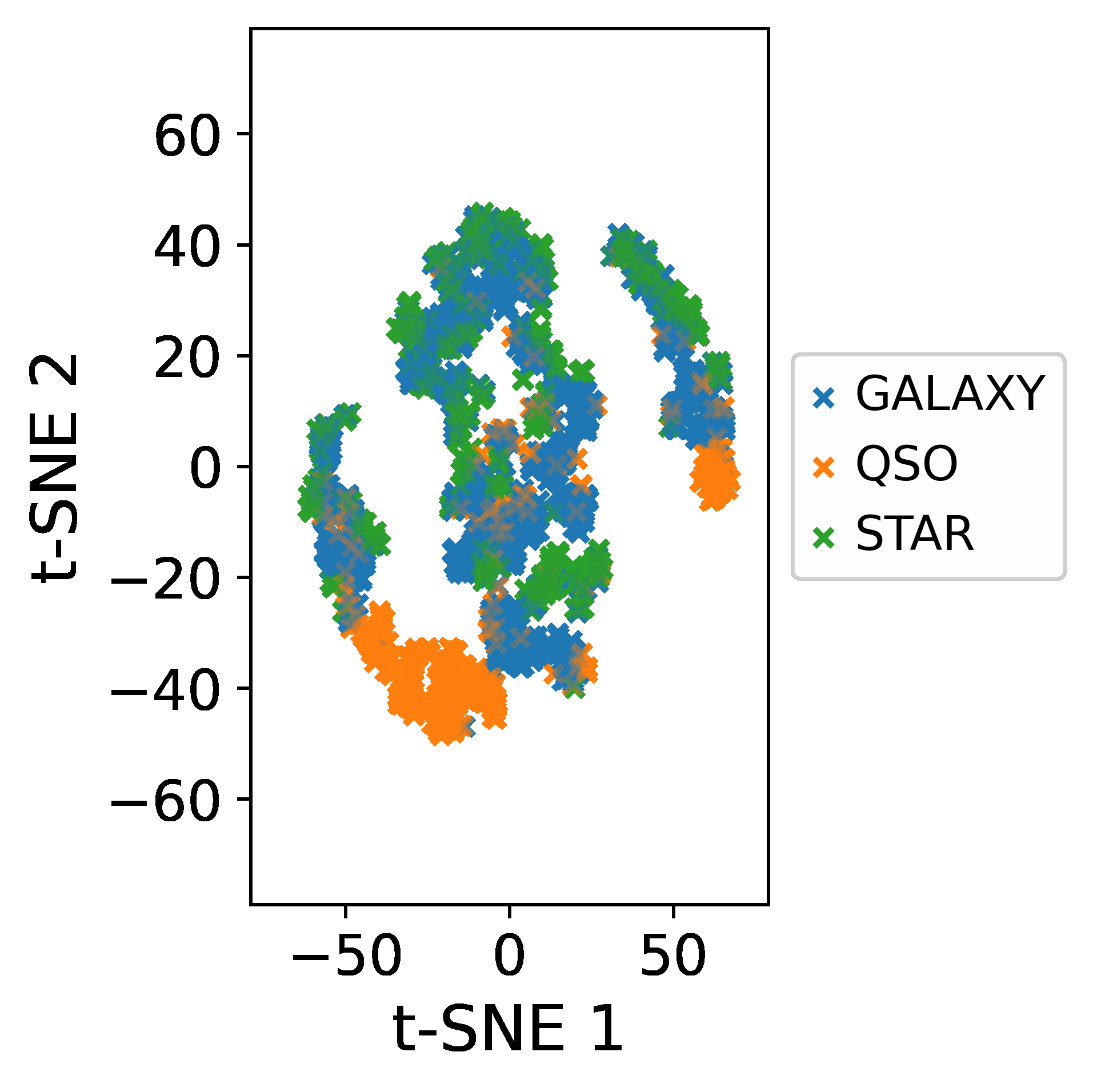

### Supplementary_Video4.gif

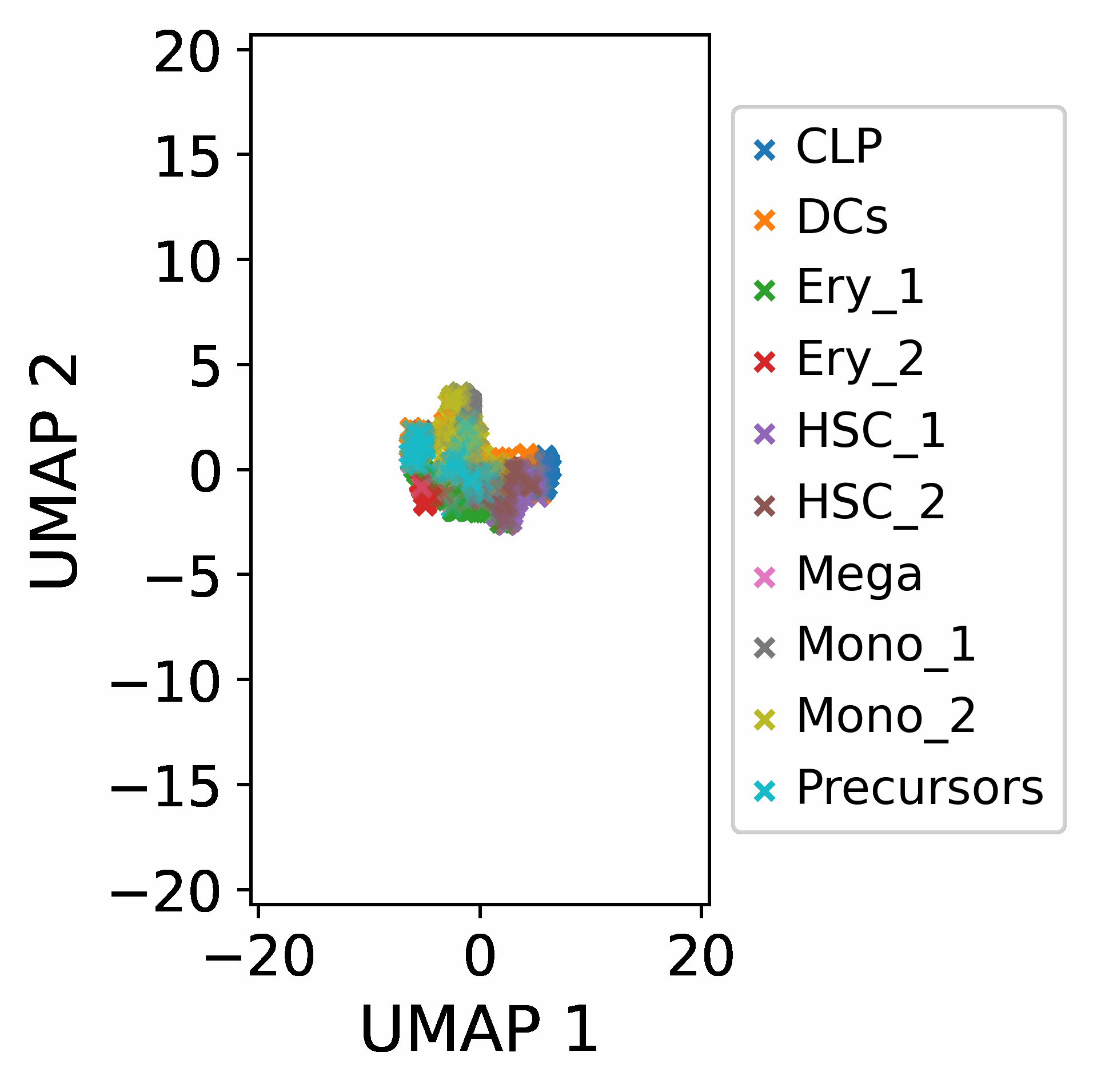
